# Mitogenomics and mitochondrial gene phylogeny decipher the evolution of Saccharomycotina yeasts

**DOI:** 10.1101/2021.06.11.448017

**Authors:** Anastasia C. Christinaki, Spyros G. Kanellopoulos, Alexandra M. Kortsinoglou, Bart Theelen, Teun Boekhout, Vassili N. Kouvelis

## Abstract

Saccharomycotina yeasts contain diverse clades within the kingdom of Fungi and are important to human everyday life. This work investigates the evolutionary relationships among these yeasts from a mitochondrial (mt) genomic perspective. A comparative study of 141 yeast mt genomes representing all major phylogenetic lineages of Saccharomycotina was performed, including genome size and content variability, intron and intergenic regions’ diversity, genetic code alterations and syntenic variation. Findings from this study suggest that mt genome size diversity is the result of a ceaseless random process mainly based on genetic recombination and intron mobility. Gene order analysis revealed conserved syntenic units and many occurring rearrangements, which can be correlated with major evolutionary events as shown by the phylogenetic analysis of the concatenated mt protein matrix. For the first time, molecular dating indicated a slower mt genome divergence rate in the early stages of yeast evolution, in contrast with a faster rate in the late evolutionary stages, compared to their nuclear time divergence. Genetic code reassignments of mt genomes are a perpetual process happening in many different parallel evolutionary steps throughout Saccharomycotina evolution. Overall, this work shows that phylogenetic studies that employ the mt genome of yeasts highlight major evolutionary events.

## 1. Introduction

Ascomycetous yeasts comprise some of the most economically and clinically important species within the fungal kingdom, since they influence many aspects of humans’ everyday life. For instance, the production of some of the most common products in human diet, like bread and wine, involves *Saccharomyces cerevisiae* (Botstein and Fink, 2011; Peter et al., 2018), while other yeasts contribute to the fermentation and flavoring of alcoholic beverages (Eldarov et al., 2016). Other members of the Saccharomycotina subphylum, like some *Candida* species, cause serious human infections (Wang et al., 2018; Pote et al., 2020). Besides their importance to humans, Saccharomycotina yeasts present several other characteristics, like single cell morphology, asexual reproduction by budding and sexual, by the formation of ascospores, as well as facultative respiration. These characteristics render them highly interesting for research, since they can be employed as single cell eukaryotic models, with *S*. *cerevisiae* as the most commonly used one (Stajich et al., 2009; Kurtzman and Boekhout 2017; Nielsen, 2019).

According to current taxonomy, Saccharomycotina consists of a single order, Saccharomycetales, in contrast with other fungal subphyla that contain several discernible orders. The main reason for this taxonomy is the ‘simple’ morphology of the budding yeasts cells, which does not provide evidence for sufficient morphological discrimination (Knop, 2011). Molecular based phylogenies classify yeasts at the family level. Recent phylogenetic studies divide this subphylum into 12 families (Shen et al., 2018), in contrast to previous studies which were based on few nuclear molecular markers that assigned them into 11 different families (Kurtzman, 2011; Kurtzman and Boekhout, 2017). Numerous studies during the last two decades showed that Saccharomycotina present high levels of genetic variability (Benítez et al., 1996; Braun, 2003; Fernández-Espinar et al., 2003; Dujon, 2006). Contrasting morphological similarity between species like *Yarrowia lipolytica* and *Saccharomyces cerevisiae*, their genetic differentiation has reached extensive levels, similar to that between roundworm and human genome (Dujon, 2006; Shen et al., 2018). This genetic diversity is attributed to several evolutionary events that took place within this lineage, like the alteration of their genetic code and the Whole Genome Duplication (WGD) events (Kellis et al., 2004; Krassowski et al., 2017). In detail, genetic code alteration occurred in three independent evolutionary events, during which the CUG codon translated Serine (two different events) or Alanine (third event), instead of Leucine (universal genetic code). As a result, phylogenetic trees support the existence of the CUG-Ser1, CUG-Ser2 and CUG-Ala distinct groups (Santos et al., 1997; Miranda et al., 2006; Krassowski et al., 2017). The second evolutionary landmark which influenced genetic diversity in yeasts is the WGD event that happened ~100 Million Years Ago (MYA) in Saccharomycetaceae (Wolfe and Shields, 1997; Kellis et al., 2004). It is proposed that WGD is the result of autopolyploidization or hybridization. However, a hybridization model was recently proposed to explain this WGD (Marcet-Houben and Gabaldón 2015), since genetic rearrangements, gene losses and mutations following the hybridization WGD event lead to genome evolution and adaptation (Wolfe, 2015; Fisher et al., 2018; Escalera-Fanjul et al., 2019).

The mitochondrion is a vital organelle due to its role in energy production of eukaryotic cells (Hatefi, 1985). Multiple mitochondria occur inside a cell which in turn contain their own genome (mtDNA) in multiple replicates, with genes encoding, among other products, proteins necessary for oxidative phosphorylation (Zamaroczy and Bernardi, 1985; Burger et al., 2003). The first sequenced mtDNAs of the *Saccharomycotina* subphylum were those of *Wickerhamomyces canadensis* and *Saccharomyces cerevisiae* (Sekito et al., 1995; Foury et al., 1998). The mt genes located in genomes of species belonging to Saccharomycotina are the typical genes found in almost all fungi and metazoa, i.e., subunits of the ATP synthase complex (*atp*6*, atp*8 and *atp*9), of apocytochrome b (*cob*), of cytochrome *c* oxidase complex (*cox*1-3), of NADH dehydrogenase (*nad*1-6*, nad*4L), the large and small rRNA (*rns* and *rnl*) and approximately 25 tRNAs (Kouvelis et al., 2004; Solieri, 2010; Dujon, 2010; Korovesi et al., 2018). Interestingly, mt genes encoding subunits of NADH dehydrogenase are absent in specific Saccharomycotina mt genomes, i.e., species belonging to families Saccharomycetaceae and Saccharomycodaceae (Pramateftaki et al., 2006; Freel et al., 2015).

The mtDNA has been utilized for comparative genomics of the phylum Ascomycota at various taxonomic levels (Freel et al., 2015; Wolters et al., 2015), as it has a different evolution rate compared to the nuclear genome, which makes mtDNA an alternative valuable source of phylogenetic markers as previously shown (Ballard and Whitlock; 2004; Kouvelis et al., 2008; Duo et al., 2012; Kortsinoglou et al., 2019). The size of the mtDNA varies greatly among Saccharomycotina species, i.e., from 18,884bp (*Hansenianspora uvarum*) to 107,123bp (*Nakaseomyces bacillisporus*) (Pramateftaki et al., 2006; Bouchier et al., 2009). The majority of yeast mt genomes are reported as circular molecules (Williamson, 2002). *Candida parapsilosis* (Kovac et al., 1984) and *Hansenula mrakii* (Wesolowski et al., 1981) were the two first described yeast species presenting linear mtDNAs. Recent studies suggest that mt genomes of yeast are linear or circular mapping, or interconvert between the two topologies (Nosek et al., 1998, Nosek and Tomáska, 2003; Chen et al., 2018). Moreover, the mt gene order (synteny) is diverse among taxa, even withing the same genus (Fricova et al., 2010; Bartelli et al., 2013; Freel et al., 2015). It has been observed that certain gene clusters are formed, which remain conserved despite the general occurring re-locations of mt regions (Pantou et al., 2008). These existing patterns help to identify gene rearrangements, partial genome inversions and duplications. The above characteristics, as well as frequent intra- and inter- species recombination of mtDNA (Fritsch et al., 2014; Brankovics et al., 2017) render mtDNA a promising marker for resolving phylogenetic relationships of Saccharomycotina yeasts.

This work aims to study the evolutionary relationships of ascomycetous yeasts from an mt genomic perspective. In detail, this work investigates the genetic elements (i.e., introns, gene clusters, intergenic regions, GC islands), genetic mechanisms (i.e., recombination) and evolutionary events (i.e., genetic code changes) under the prism of molecular dating. Moreover, it studies their impact on shaping the existing phylogenetic relationships within this subphylum. For simplicity reasons, in the rest of the following manuscript the term “yeasts” will refer to yeasts of Saccharomycotina, unless stated otherwise.

## 2. Materials and methods

### 2.1 Retrieval of DNA sequences

In this work, 151 mt genome sequences were examined. Out of the 151 species, 141 belong to the subphylum of Saccharomycotina and 10 belong to the subphylum of Taphrinomycotina and were used as outgroups. The 141 yeast species belong to 11 out of the 12 Saccharomycotina families, since representative genomes for the 12^th^ family were not available when the analyses were performed (last search 01-December-2020). The majority of the sequences (131 out of the 151), including the species used as outgroups, were retrieved from complete mt sequence entries of GenBank. Eleven taxa were collected from various databanks, but a complete mt genome was not found and as a result, the precise gene limits for these taxa were not detected. For this reason, protein sequences were identified and retrieved using protein similarity searches with BLASTp (Altschul et al., 1990) and were further used in phylogenetic analyses (Table 1, Table S1). Moreover, mt genomes of 10 taxa were extracted from whole genome sequencing (WGS) projects (Table S2), by using the tBLASTn/BLASTx/BLASTn tool for each WGS scaffold and comparing them with mt genome sequences of closely related species. In cases where mt genomes belonged to more than one scaffold, different parts were assembled into one sequence, using the Seqman tool of Lasergene Suite 11 (DNASTAR Inc., Madison, WI) (Burland, 2000).

**Table 1:** The families of Saccharomycotina examined in this study. The number of species/strains from each family is provided (details in Table S1)

| Family | Number of taxa | Phylogenetic group (clade) | Number of taxa |
| --- | --- | --- | --- |
| Phaffomycetaceae | 9 | Phaffomycetaceae | 9 |
| Debaryomycetaceae | 37 | CUG-Ala | 3 |
| Metschnikowiaceae | 4 | CUG-Ser1 | 40 |
| Saccharomycopsidaceae | 1 | CUG-Ser2 | 1 |
| Trichomonascaceae<br>(/Dipodascaceae) | 2 | Dipodascaceae/<br>Trichomoscaceae | 14 |
| Dipodascaceae | 11 | Lipomycetaceae | 3 |
| Trichomonascaceae | 2 | Pichiaceae | 11 |
| Lipomycetaceae | 3 | Saccharomycetaceae LKE | 16 |
| Pichiaceae | 12 | Saccharomycetaceae post-WGD | 27 |
| Saccharomycetaceae | 55 | Saccharomycetaceae TZ | 11 |
| Saccharomycodaceae | 2 | Saccharomycodaceae | 2 |
| Trigonopsidaceae | 3 | Sporopachydermia | 1 |
|  |  | Trigonopsidaceae | 3 |
| Total | 141 | Total | 141 |

### 2.2 Genome Annotation

To proceed with the mt genome analyses, annotation of all the acquired genomes was verified manually. The protein coding and the ribosomal (rRNA) genes were identified using BLASTx and BLASTn (Altschul et al., 1990), respectively, after comparisons with known related sequences. The *trn* genes were detected using the online software tRNAScan-SE (Chan and Lowe, 2019). The mt genomes of 32 species were re-annotated (Table S2), since mis-annotations regarding gene limits and intron content were detected. Moreover, 9 mt genomes were *de novo* annotated along with 10 genomes found in databanks as WGS projects (Table S2).

### 2.3 Genome analyses

Following the correction of genome annotations, comparative gene order (synteny) analyses were performed manually to locate any existing differentiations or similarities in genome organization among taxa belonging to different phylogenetic groups. To further extend our analyses, the total number of mt genes in each genome was recorded to detect any possible deletions or duplications, mostly of *trn* genes. Our study included genome size comparisons, as well as an analysis of intron abundance and their distribution among all genes. Intron characterization was performed using RNAweasel and MFannot (Lang et al., 2007).

### 2.4 Codon usage frequency

Alternative codon usage frequency was examined in all taxa included in this study. Fungal mt genomes utilize both alternative codes to the Universal Genetic Code (NCBI code 1), i.e., NCBI genetic code 3 (The Yeast mitochondrial genetic code) and 4 (The Mold, Protozoan, and Coelenterate Mitochondrial Code and the Mycoplama/Spiroplasma Code) and thus, they present a few alternative codons encoding for the amino acids Threonine (T), Methionine (M) and Tryptophan (W). Codon usage frequency was examined using Sequence Manipulation Suite (SMS) (Stothard, 2000), after inserting the gene sequence and the genetic code used by the species’ mt genome. The related information regarding the mt genetic code utilized by each species was retrieved from the NCBI Taxonomy database. In case where codon frequency results were contradictory and unusually different from the rest, the mt genetic code was further examined and verified or even corrected in some species. Seven out of the 14 protein coding genes were included in this study, since seven genes of NADH dehydrogenase subunits 1-6 and *nad*4L are not present in all studied fungal genomes, and thus would provide misleading evidence regarding comparative codon usage frequency of the whole mt genome. After listing the frequency of alternative codons, the RSCU index was calculated as an indication of possible codon usage bias (Sharp et al., 1986). The RSCU value for a codon reflects the observed frequency of that codon, divided by the expected frequency, assuming an equal usage of the synonymous codons for an amino acid (Sharp et al., 1986). The synonymous codons with RSCU values > 1.0 have positive codon usage bias and are defined as abundant codons, whereas those with RSCU values < 1.0 have negative codon usage bias and are defined as less-abundant codons.

In detail, the frequencies of synonymous codons within each amino acid class were calculated utilizing mt sequences of the protein coding genes *cob*, *atp*6*, atp*8*, atp*9 and *cox*1-3. In the above analyses, codon frequencies and RSCU values were measured for each taxon and for each of the above genes. In addition, accumulative mean RSCU values were measured for a) each codon and each gene separately, by adding the respective values of all taxa (e.g. ATG of *cob* in all species) and dividing them with the total number of taxa (Table S3), b) each codon in all genes of each taxon (e.g ATG of *cob* plus ATG of *cox*1 of each strain etc.) divided with the number of genes (Table S3) and c) a final mean RSCU value was calculated for each codon by summing RSCU values occurring in calculation and dividing them with the total number of taxa (Table S3). In all cases, RSCU values were measured separately for a) species utilizing genetic code 3 and 4, and b) for each codon within each family or phylogenetic cluster, respectively.

### 2.5 GC cluster analyses

To investigate the role of concatenated GC nucleotides, all repeats present in the mt genomes of organisms belonging to the Saccharomycotina subphylum were initially detected using the UGene program (Okonechnikov et al., 2012), for each species separately. The minimum length of a GC nucleotide repeat was set to 30 bp and the maximum distance among repeats was increased to 20000 bp compared to the 5000 bp of the default parameters, in order to expand our search (more “relaxed” conditions). Repeat analyses were performed at 90% identity for the purposes of determining repeats. All other parameters were set to default. The repeats that contained less than 50% GC were discarded and then the repeats of each organism were clustered with CD-hit-est (Ying et al., 2010) with identity set to 80% and all other parameters at default. The representative sequence of each cluster was selected, and an alignment was performed with the representative sequence of all species using Clustal Omega (Madeira et al., 2019) and the consensus of this alignment was used to create DNA logos with WebLogo (Crooks et al., 2004).

### 2.6 Phylogenetic analyses

Amino acid sequences of all 14 protein coding genes usually found in mt genomes, i.e., Apocytochrome b (*cob*), Cytochrome c oxidases (*cox*) subunits 1-3, ATPase subunit 6,8,9 and NADH dehydrogenases (*nad*) subunits 1-6, *nad*4L, were collected in order to construct the concatenated matrix. The collected amino acid sequences were aligned using ClustalW (Thompson et al., 1994) as implemented in Lasergene’s MegAlign v.11 program (Burland, 2000). Alignment parameters were set to default and the result was verified manually in order to ensure correct alignments. A concatenated matrix for all 14 protein coding genes was created for the phylogenetic analyses. In species lacking the *nad* genes, the corresponding sites were replaced with the symbol of missing information.

Phylogenetic trees were constructed using Neighbour Joining (NJ), Bayesian Interference (BI) and Maximum Likelihood methods (ML), using PAUP (Wilgenbusch and Swofford, 2003), MrBayes (ver. 3.2) (Ronquist and Huelsenbeck, 2003) and RAxML (Stamatakis, 2014), respectively. 7 *Pneumonocystis* species and 3 *Schizosaccharomyces* species were used as outgroups. The methodology applied for NJ, BI and ML phylogenies has already been described in previous studies (Korovesi et al., 2018; Kortsinoglou et al., 2019). In detail, for the NJ analyses, reliability of nodes was assessed using 1M bootstrap iterations for all the 14 individual and concatenated datasets. For the BI analyses the program ProtTest (ver. 3) (Abascal et al., 2005) was used in order to determine the model that was best suitable for our dataset. In the case of the concatenated dataset, the Bayesian Information Criterion (BIC) was applied, and the best amino acid substitution model was found to be mtREV, with Gamma Distributed (G) rates among sites. Four independent MCMCMC searches were performed, using 10M generations and sampling set adjusted every 100,000 generations. Convergence was checked visually by plotting likelihood scores vs. generation for the two runs. Burn-in was set to default in all cases.

### 2.7 Molecular dating

To estimate the divergence times among the subphylum Saccharomycotina, the concatenated tree, as produced from the Bayesian analysis, was used as input in the RelTime method of MEGA7 (Kumar et al., 2016). Four calibration points were used for the analysis as follows: (1) *Saccharomyces cerevisiae – Saccharomyces uvarum* split (14.3-17.94 MYA), (2) *Saccharomyces cerevisiae – Kluyveromyces lactis* split (103-106 MYA), (3) *Saccharomyces cerevisiae – Candida albicans* split (161-447 MYA), (4) Saccharomycotina – Taphrinomycotina (304-590 MYA) (Shen et al., 2018; Kumar et al, 2017). For the analysis, the mtREV amino acid substitution model with Gamma Distributed (G) rates among sites (α = 0.6990) was used. The estimated log likelihood value was −291342.08.

## 3. Results

### 3.1 General characteristics of mt genomes

The 141 yeast mt genomes (Table 1) studied in this work presented a few similar characteristics and several different properties (Table S1). The size of the mtDNA ranged from 18,884 to 107,123 kb with an average size of 43,476 kb. The size of most genomes was distributed between 20-39.9 kb, representing 53.8% of the total examined mt genomes or 70.7%, if mt genomes with 40 to 40.9 kb were included (Table S1). In all fungal mt genomes, the gene content of protein coding genes related to oxidative phosphorylation and production of ATP was conserved. As an exception, the genes of the *NADH* complex were absent in the families Saccharomycetaceae and Saccharomycodaceae (Table S1).

The majority of yeast families presented a uniform pattern of presence or absence of *rps*3, a gene that contributes to the assembly of the small ribosomal subunit. Specifically, this gene was present in members of the families Phaffomycetaceae, Dipodascaceae, Pichiaceae, Saccharomycetaceae and Trichomonascaceae, while it was absent in all other yeast families. One exception was found in the Saccharomycodaceae family represented here by two species, i.e., *Saccharomyces ludwidgii* and *Hanseniaspora uvarum*, in which the gene *rps*3 was present in the first and absent in the latter (Table S1). The number of *trn* genes fluctuated among 21-31, with an average of 24 (Table S1).

Gene content of yeast mt genomes was mostly conserved and if introns were not considered, all mt genomes were approximately of the same genome size. The mean size of all 14 mt genes combined was 16,969 bp (Table S1). The genome size variability observed was explained by the diversity of introns and intergenic regions (Table S1). For instance, the examined mt genome of *N. bacillisporus* (107,123 kb) presented intergenic regions equaling 88% of the entire genome and no intronic sequences. The genome with the smallest proportion of intergenic regions was the one of *Millerozyma farinosa,* since only 7% was attributed to intergenic regions, while 48% of its genome size corresponded to introns (Table S1).

Yeasts genomes were either linear (19 species examined) or circular mapped (112 species examined) (Table S1). It was found that both size and genome structure variation occur scattered among all yeast families without a clear pattern. However, the general size tendency of mt genomes was clearly towards 20-50 kb, and the majority of them were circular mapped.

### 3.2 Mt concatenated gene phylogeny

The sequences of the 14 conserved mitochondrial proteins were used to determine the phylogenetic relationships of yeasts. Single gene phylogenies presented contradictory results (File S1), and thus, a concatenated approach was followed. The tree produced by Bayesian analysis was well supported, with almost all posterior probability (PP) values at 100% (Fig. 1). This result was mostly in accordance with the trees produced by the NJ and ML methods. Differences were found in some underrepresented families (Fig. 1). In the tree produced by this work, the 141 yeast taxa formed 11 phylogenetic clades with almost identical topologies to those obtained by the nuclear based phylogeny (Shen et al., 2018), with only the following differences in the mt phylogeny: (a) the Trigonopsidaceae family formed an earlier diverging lineage than Lipomycetaceae, (b) *H. uvarum* was placed basally to the CUG-Ser1 clade apart of *S. ludwigii*, the other member of the family, (c) *Komagataella pastoris* appeared to be inside the CUG-Ala clade and not in Pichiaceae, its closely related family, (d) the only available mt genome of *Saccharomycopsis malanga* (representing CUG-Ser2 clade) was basal to the clade containing the families Saccharomycetaceae and Phaffomycetaceae and it did not seem to be related to the CUG-Ser1 clade (Fig. 1 and Shen et al., 2018).

**Fig 1:**
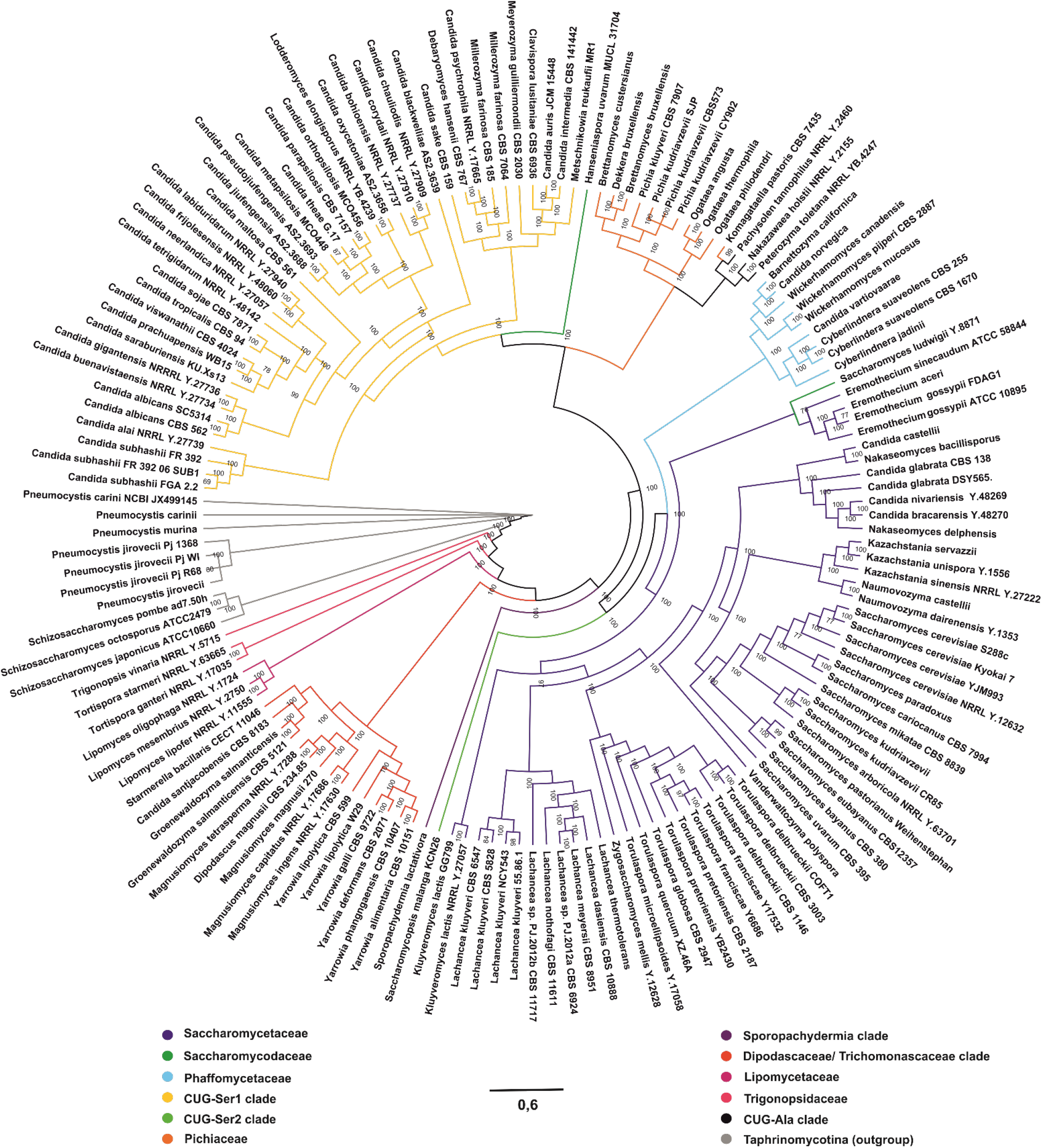
The phylogenetic tree of all yeast species based on the concatenated dataset of the 14 conserved mt proteins (i.e., Atp6,8–9, Cob, Cox1-3, Nad1-6 and 4L) as produced by the Bayesian Inference (BI) method. All topologies produced are in accordance with the respective NJ and ML methods. The species belonging to Taphrinomycotina subphylum were used to root the tree. The clades are colored according to taxonomic families as follows: Saccharomycetaceae blue, Saccharomycodaceae dark green, Phaffomycetaceae light blue, GUG-Ser1 clade yellow, CUG-Ser2 clade light green, Pichiaceae orange, Sporopachydermia clade dark purple, Dipodascaceae/ Trichomonascaceae clade red, Lipomycetaceae light purple, Trigonospidaceae pink, CUG-Ala clade black and Taphrinomycotina grey (outgroup).

### 3.3 Molecular dating

Molecular dating was used to estimate the divergence of the Saccharomycotina subphylum as presented by the mitochondrial phylogeny (Fig 2, Fig S1 and Table S4). According to the mt based phylogeny, (a) the subphylum originated approximately 590 MYA, (b) the WGD group diverged 75.32 MYA from the ZT clade (Zygosaccharomyces/Torulaspora) of the Saccharomycetaceae, (c) the two CUG-Ser clades diverged independently from their closer relatives at 325 and 310 MYA, respectively and, (d) the CUG-Ala clade diverged from Pichiaceae at 181 MYA (Fig 2).

**Fig 2:**
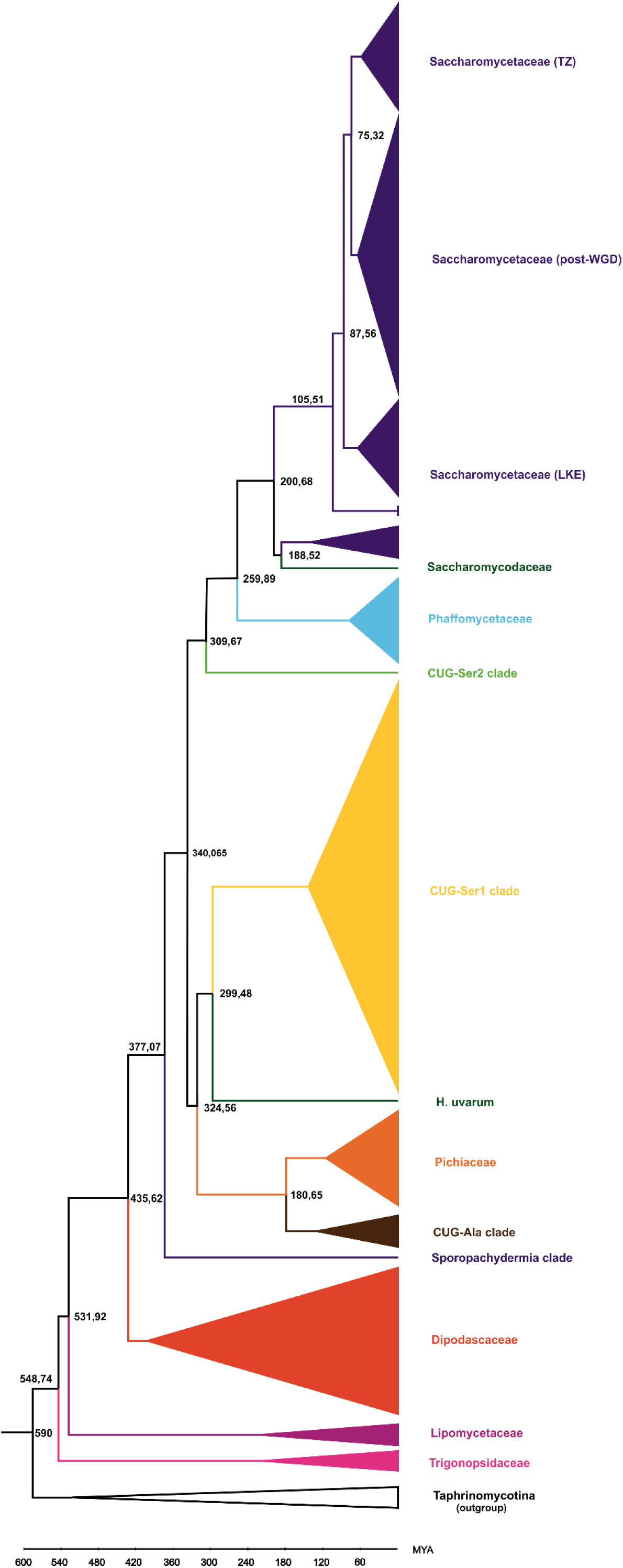
Time-Calibrated Phylogeny of the Saccharomycotina subphylum. The 141 mt genomes analyzed in this study are shown in clusters according to the families/ phylogenetic clades of the tree shown in Fig 1. For the most important nodes of the tree, the time calibration is shown in MYA. The clades are colored according to taxonomic families as follows: Saccharomycetaceae blue, Saccharomycodaceae dark green, Phaffomycetaceae light blue, GUG-Ser1 clade yellow, CUG-Ser2 clade light green, Pichiaceae orange, Sporopachydermia clade dark purple, Dipodascaceae/ Trichomonascaceae clade red, Lipomycetaceae light purple, Trigonospidaceae pink, CUG-Ala clade brown and Taphrinomycotina black (outgroup).

### 3.4 Synteny

Gene order analysis revealed a clear pattern of genome organization among all taxa studied. This pattern was based on the existence of several gene clusters consisting of both *trn* genes, and protein or ribosomal coding genes. These genes were considered to form clusters, because they were found intact (at the same or inverted orientation) in different locations of mt genomes of various species. Based on the presence of these conserved clusters, five major groups and several sub-groups of taxa could be determined (Fig 3, Table S5 and Table 2). The only criterion for grouping various taxa in one main group was the existence of at least two identical or highly similar syntenic groups in all its members (Table 2 and Table S5). Therefore, Group A contained species belonging to clades Saccharomycetaceae post-WGD, TZ, LKE and Phaffomycetaceae, and Group B consisted of the Metschnikowiaceae and Debaryomycetaceae families. Group C corresponded to species of Pichiaceae, while species that belong to the CUG-Ala clade comprised Group D. Finally, species belonging to the Dipodascaceae and Trichomonascaceae families represented group E (Fig 3 and Table S5). Species belonging to the Lipomycetaceae and Trigonopsidaceae clades were underrepresented and did not present enough similarities with the above groups. For this reason, it seemed prudent that these groups may be considered as separate additional groups which, however, need further examination by adding sequence data from more species.

**Table 2:** Groups of taxa (A-E) and the syntenic gene clusters (1-7) formed by comparing the genome organization of species examined in this study. *except for Kluvyeromyces spp. and Eremothecium spp. and Phaffomycetaceae family, **except for C. buenavistaensis NRRL Y-27734, C. albicans SC5314 and CBS 562, ***(only in C. galli and Yarrowia spp.)

|  | Group A | Group B | Group C | Group D | Group E |
| --- | --- | --- | --- | --- | --- |
| 1 | *trnF*-(*trnC*)-(*trnV*)* | *atp6-atp8-trnP-trnC-trnG* | *rps3-trnP-trnR-trnV* | *nad6-nad1-nad4* | *cob-nad2-nad3* |
| 2 | (*trnL*)-*trnQ-trnK*-(*trnR*) | *trnL-trnY-trnH*-(*trnM*)-*trnT-trnE* | (*cob*)-*nad2-nad3* | *cox2-trnS-trnA* | *atp9-cox2-nad4L-nad5\*\*\** |
| 3 | *trnG-trnD-trnS*<br>*trnG-trnD-trnS-trnS/trnR*<br>*trnR-trnG-trnS*-(*trnS*) | (*trnR*)-*trnV-trnW-trnF*-(*trnK*-*cox3*)<br>*trnR-trnV-trnW-trnF*-(*trnR*) | (*nad4*)-(*trnR*)-*trnS-trnD-trnA* | *nad2-nad3-cox1* | (*trnT*)-*trnE-trnM-trnM-trnL-trnQ-trnK* |
| 4 | *trnY-trnN-trnA-trnI*-(*trnS*)<br>*trnA-trnI-trnY-trnN*<br>*trnI-trnS-trnA-trnY-trnN*<br>*trnI-trnA-trnN-trnY* | *trnQ-trnD-trnS*-(*trnR*) | (*trnL*)-*trnQ-trnK-trnH*-(*trnM*) |  |  |
| 5 |  | *trnS-trnL-nad4* |  |  |  |
| 6 |  | *cox2-trnN-nad6-nad1* |  |  |  |
| 7 |  | (*trnM*)- *rns-trnI* **<br>*trnM<sub>1</sub>-trnM<sub>2</sub>-rns*<br>*trnM<sub>1</sub>-trnM<sub>2</sub>-rns-trnI* |  |  |  |

**Fig 3:**
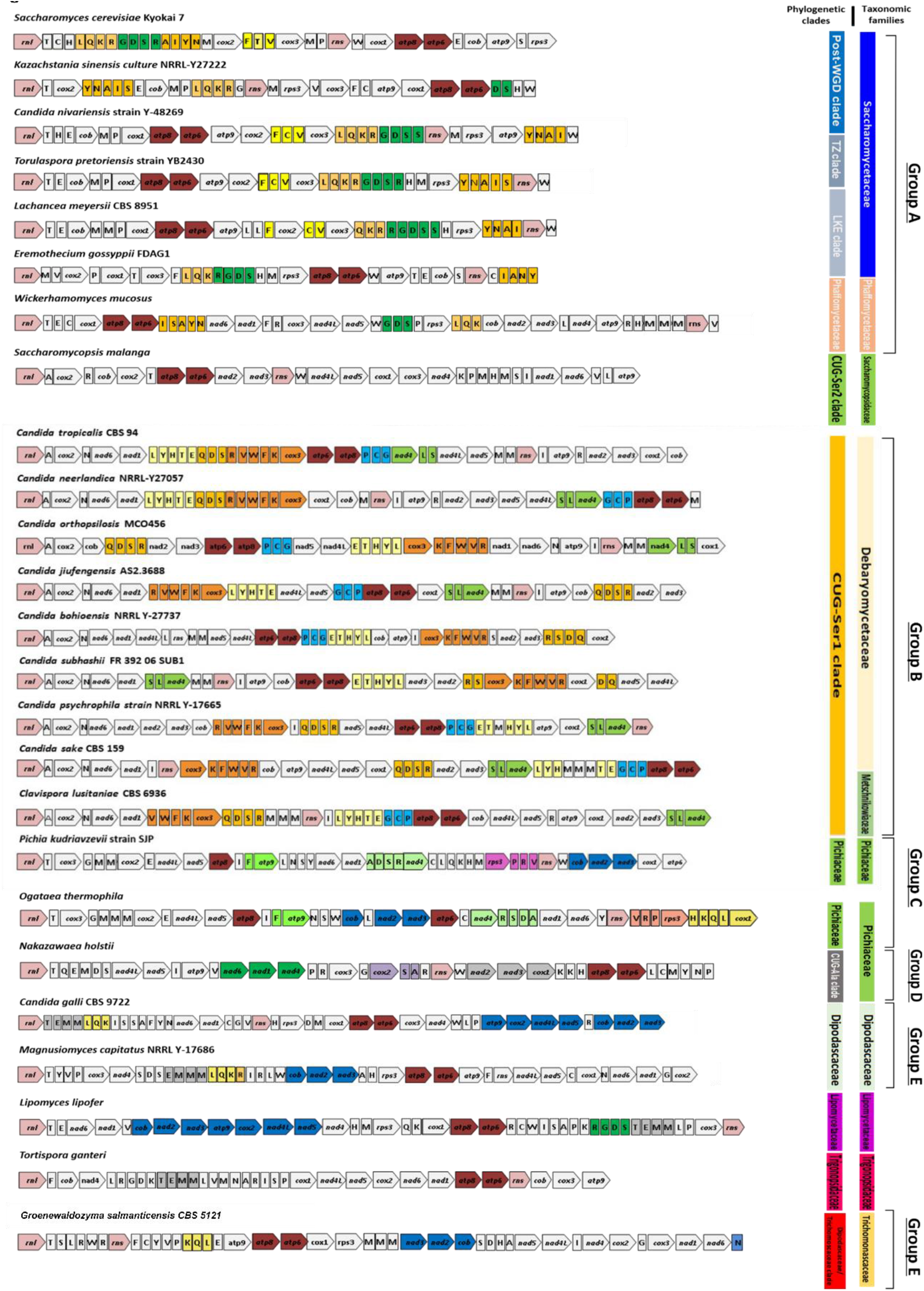
Synteny of mt genomes for representative species of all families/ phylogenetic clades. Genes are shown as boxed arrows and conserved syntenic units are highlighted with different colors.

Taxa belonging to Group A had a small degree of intra-species rearrangements in all of the subgroups (Fig 3 and Table S5). It was possible to distinguish four main gene clusters (Table 2). Despite the conserved anchoring of these clusters at intra species level, there was a high rate of gene rearrangements inside each individual cluster. The gene content was the same, but the gene order was different in each syntenic group. Characteristic examples were the rearrangements of syntenic groups *trn*G-*trn*D-*trn*S-*trn*R and *trn*Y-*trn*N-*trn*A-*trn*I (see Table 2 and Table S5 for alternative orders).

As previously mentioned, taxa belonging to Group B had seven main syntenic groups (Fig 3, Table 2, Table S5). These maintain a conserved gene content but were subjected into several major whole cluster rearrangements. As a result, this gene shuffling created newly reversed, joined or divided clusters. Taxa belonging to groups C, D and E presented smaller syntenic units of usually two to three genes (Fig 3, Table 2, Table S5).

### 3.5 Genetic codes and codon frequency

The study of 141 strains belonging to the Saccharomycotina subphylum showed that the species belonging to families Debaryomyceteceae, Metschnikowiaceae (i.e., both comprising phylogenetic clade CUG-Ser1), Dipodascaceae, Lipomycetaceae, Pichiaceae, Phaffomycetaceae, Trichomonascaceae, Trigonopsidaceae and the CUG-Ala phylogenetic clade employ genetic code 4 for translating their mt genes, while almost all species of Saccharomycetaceae (i.e., LKE, TZ and post-WGD clades with the exception of *Kluyveromyces* spp) and *Saccharomyces ludwigii*, *H. uvarum* and *Pachysolen tannophilus* use genetic code 3 (Tables S1 and S3).

Results showed that accumulative Relative Synonymous Codon Usage (RSCU) values of individual genes, as well as the overall RSCU values presented highly biased frequencies for ATG (Met), TGA (Trp), ACA and ACT (Thr), with the exception of genes *atp*8 and *atp*9 in the TGA codon (Fig 4 and Table S3). The ATA codon was used by species utilizing genetic code 3 with an approximate ratio of 1:5 to the synonymous ATG codon. In a similar way, the TGG, which is the universal codon for tryptophan, was mainly found in yeast genomes utilizing genetic code 4, in a ratio of 1:5 to the respective TGA codon. On the contrary, species using genetic code 3 had a higher percentage of the TGA codon (98.2%) compared to the TGG (1.8%) (Fig 4). In all taxa studied, the most abundant Thr codons were ACA and ACT (Fig 4). As expected, CTN (Thr) codons were only detected in species using genetic code 3, and the most abundant codons, after ACA and ACT, were CTA and CTT (Fig 4 and Table S3). Trp codons were not detected in *atp*9, while in *atp*8 these codons were absent only in species utilizing genetic code 4 (Table S3). The gene *atp*9 had a high number of ACA codons in taxa belonging to both genetic codes.

**Fig 4:**
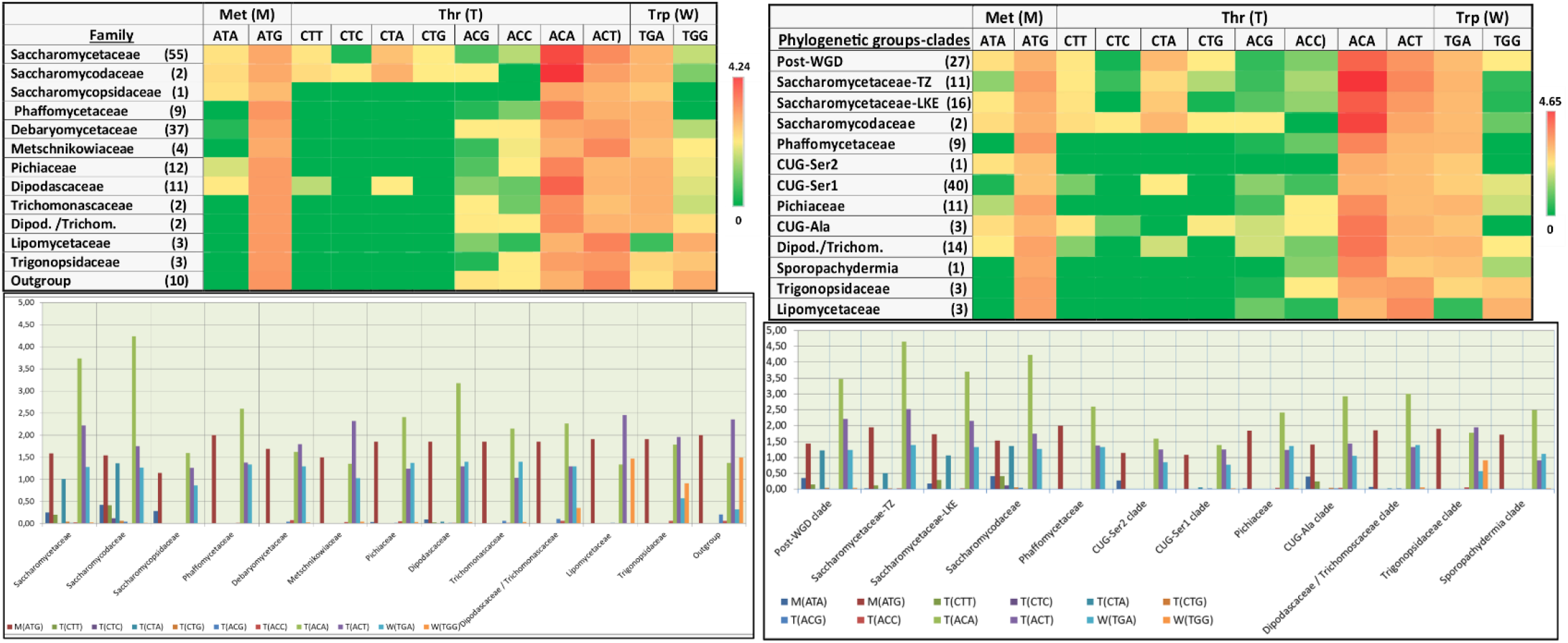
Mean RSCU values (heatmaps) for each codon of a) all families and b) all phylogenetic groups. Mean RSCU values were calculated using the respective mean values of each genome, from all taxa belonging to each clade/phylogenetic group.

### 3.6 Intron distribution

The mt genes of the examined yeasts presented an interesting intronic distribution concerning intron types and their gene hosts. Most introns were found in the *cox*1 gene (517 introns) which usually contains IB introns (388 introns) and in a lesser extent II and ID introns (i.e., 57 and 50, respectively) (Table S6I.b). The *cob* gene was the second most intron-rich gene (193 introns), followed by *rnl* (89 introns) and *nad*5 (35 introns) (Table S6I.b). Genes *atp*6, *cox*2, *cox*3, *nad*1, *nad*4 and *rns* contained fewer introns (i.e., less than 10) compared to the above genes, while *atp*8, *atp*9, *nad*2, *nad*3*, nad*6 and *nad*4L lacked introns in the mt genomes of the yeasts examined (Table S6I.b). The most common type of intron in the *cob* gene was ID (84 ID out of 193 introns found in *cob*), followed by the IB intron (57) (Table S6I.b). The most abundant intron of *rnl* was IA (60 out of 89 introns found in *rnl*) (Table S6I.b). Only 16 group II introns were found in four different genes, i.e. *cox*1 (11 introns), *cob* (3 introns) and both *cox*2 and *nad*5 (1 intron) (Table S6II).

IB introns were the most abundant within the various mt genes of all genomes studied, regardless of the family or phylogenetic clade (Table S6I.a). In detail, in all families, IB introns represented more than 50% of the total introns found, with the lowest value (39.3%) in the case of Pichiaceae and the largest values (80% and 66.7%) in Sporopachydermia clade and Saccharomycetaceae of the LKE clade, respectively (Table S6I.a). Five phylogenetic clades, i.e., Debaryomycetaceae and Metschnikowiaceae (both comprising the CUG-Ser1 clade), Saccharomycetaceae (belonging to the TZ and post-WGD clades), Saccharomycodaceae, and Trichomonascaceae showed an IB intron distribution of approximately 50%. Introns of subtypes ID and IA were the next most commonly found ones within the yeast phylogenetic clades, with 16.28% and 12.56%, respectively (Table S6I.a). Group II introns were located in only 10 species, which either contained more than one group II introns (i.e., *S. malanga* and *S. ludwigii* all found in *cox*1) or in fungal species taxonomically distributed at CUG-Ala clade and the “early diverging” families like Lipomycetaceae, Sporopachydermia and Trigonopsidaceae (Table S6II).

When examining the intron distribution in relation to the hosting mt gene and the family or phylogenetic clade in which these genes were located, it became obvious that the most intron rich genes were those of *cox*1 of the post-WGD Saccharomycetaceae clade, followed by *cox*1 of the Dipodascaceae family and *cob* of the post-WGD Saccharomycetaceae clade, with 116, 62, and 55 cases, respectively (Table S6I.c). Introns of Dipodascaceae and Trigonopsidaceae families were distributed in seven and eight genes, respectively, while in the rest of the families, introns were only located in one to five genes (Table S6I.c).

Ιntron insertion sites were correlated with both their included ORF (Homing Endonuclease Gene - HEG or Reverse Transcriptase gene - RT) and their hosting gene and they presented three different motifs of intronic distribution: (a) introns, while carrying intronic ORF of similar type, may be found as ancestral elements at the same insertion site of a mt gene, (b) they may be mobile from one gene to another (due to similar insertion site) and (c) their intronic ORF may independently be mobile or lost (Table S6I and Table S6II). A single example which covers all three mentioned observations could be detected in species belonging to families Trigonopsidaceae, Lipomycetaceae, Dipodasceceae, Trichomonscaceae and the CUG-Ser2 clade, for which the *cob* gene contained an intron at position 429 bp. This intron belonged to subtype ID and carried a LAGLIDADG homing endonuclease gene. It was inserted at a conserved position (5’ WYWNTGRGGT ↓ GCWACWGTWA 3’) for all species examined (Table S6II). However, in three different *Yarrowia* species (*Y. deformans, Y. galli*, and *Y. lipolytica*) a similar intron ID carrying the LAGLIDADG gene was found at a similar insertion site of *cox*3 (position 444 or 528 bp of gene). Moreover, in the case of *Y. galli* strain CBS9722 the third hypothesis of this study could be confirmed, as at the same insertion position of *cob* gene an ID intron was located but it was ORF-less (Table S6II).

### 3.7 GC clusters

The analysis of GC-clusters showed that 59 of the mitogenomes did not contain any GC repeats that meet the applied search requirements (see Materials and Methods), while 48 of the species had less than 10 repeats and the remaining 34 had repeats ranging from 10 to 435 (*Yarrowia prachuapensis* WB15 presented the maximum number of 435 repeats) (Table S1). The number of GC rich repeats seemed to be positively related to the size of the mitogenome (p = 2.616e-11, rho = 0.543) according to the Spearman correlation, while it was not related to GC content (p-value=0.932, rho=−0.0076). The examination of these repeats for each species individually showed sequence conservation and revealed the formation of distinct groups in each different organism based on this sequence similarity. Representative sequences, one of each GC clusters from all species, were aligned and showed that there is a highly conserved core of GC nucleotides in most of the species that had more than 10 GC rich repeats (File S2).

## 4. Discussion

Undoubtedly, mt genomes in all yeast species exhibit a mostly conserved gene content but they present high size diversity (Kouvelis et al., 2004; Lang et al., 1999). This variability is analogous to that observed in genomes of the species belonging to Pezizomycotina, the other large subphylum of Ascomycota (Megarioti and Kouvelis, 2020). In this work, it was confirmed that the mt genome of *H. uvarum* is the smallest fungal mtDNA currently observed (Pramateftaki et al., 2006). For this species, the functional size of the mt genome is intriguingly smaller (approximately 11 kb), if the telomeric repetitions of this linear genome are excluded. The lack of mt *nad* genes may explain the small size, contrasting with the respective genomes of *S. cerevisiae*, in which *nad* genes are also absent but their mt genome sizes vary from 73,450 bp to 95,658bp (De Chiara et al., 2020). In this work, it was shown that intergenic regions and intron abundance are responsible for genome size diversity, but these are not always analogous with total genome size fluctuations (Table S1), despite what is shown in other studies (Wolters et al., 2015; De Deng et al., 2018; Chiara et al., 2020). Moreover, there is no clear pattern that indicates the relation of mt genome size with the yeasts’ phylogeny. Similarly, mt genome morphology (linear or circular mapping) is randomly found across the phylogenetic trees. Thus, both size and morphology of yeast mt genomes may change in a ceaseless random manner throughout the species phylogeny. This size variability is found throughout the subphylum of Saccharomycotina (i.e., in all families examined in this work) and thus, it seems to be in accordance to the recently proposed theory that genetic drift is the main factor for size differentiation of mt genomes among yeasts belonging to the LKE clade (Xiao et al., 2017). However, genetic drift is an incidental driven process without any clear direction, which usually occurs in small-sized populations (Masel, 2011). Therefore, while genetic drift cannot be precluded as a possible mechanism of mt genome diversity, in this study it is shown that genetic hitchhiking (or genetic draft) may be considered as another driving force (Hill, 2020). Genetic hitchhiking entails natural selection and is most probably the result of on genetic recombination (Smith and Haigh, 1974). However, it is not evident that natural selection in yeasts may have played such a role in the observed mt genome diversity, since several “back and forth” mt genome changes have been observed throughout the evolution of all yeast species in this study. Nevertheless, the influence of the external environment in shaping mt genomes through directional selection has been previously shown for the animals (Ballard and Pichaud, 2014; Barreto et al., 2018). The arbitrary but perpetual changes found in yeast mt genomes, seem similar to the proposed mechanism of selective sweeps that was proposed for mammals (Meiklejohn et al., 2007). Still, mt genetic recombination is the key mechanism for shaping mt genome diversity. Irrelevant to its type (i.e., homologous, non-homologous, site-specific recombination and transposition), genetic recombination may act as a solution to the cell’s needs for genome repair to assemble a functional electron transport chain (Stein and Sia, 2017). Recombination requires repetitive identical sequences of variable size (depending on the type) and its results are usually the duplication or elimination of coding and non-coding regions. In favor of this argument, it was shown that the GC repeats presented a highly conserved core of GC nucleotides which is often reversed and complementary (File S2). This may be the cause of many recombination events (Dieckmann and Gandy,1987) as most of these GC clusters have been reported to be located in intergenic regions (Zamaroczy and Bernardi, 1986). Moreover, the existence of *trn* genes at the bilateral ends of the described syntenic units (Fig 3 and Table S5) may contribute to genetic recombination. These *trn* genes at the boundaries of the syntenic units may form a potential hairpin-like secondary structure which may play the role of the repetitive elements needed for recombination, as already proposed for metazoa (Saccone et al., 1999) and Pezizomycotina (Pantou et al., 2008).

The intriguing outcome of recombination is gene shuffling. In this work, it became evident that the mt gene order of all species belonging to Saccharomycotina could split taxa into two highly and three less populated groups, respectively, each presenting a distinct clustering pattern (Fig 3, Table S5, Table 2). The correlation of the syntenic groups with the respective phylogeny of species involved, revealed that gene order not only follows, but may also affect evolution of the mt genomes. Gene clusters observed in this work were found intact in almost all syntenic groups but in different sequential order (Fig 3, Table 2). Moreover, “early diverging” yeasts, like *Tortispora* sp., *Trigonopsis* sp., and *Lipomyces* sp., do not present a significant mt genome organization, as syntenic groups are not formed among species of the same genera (Fig 3, Table S5). All the above indicate that “organized” synteny of mt genomes with certain but variable gene clusters is a later acquired characteristic during the evolution of the Saccharomycotina subphylum. Nevertheless, the presence of specific small syntenic units, like *atp*8*-atp*6, is conserved in almost all of 141 taxa examined. Similarly, conserved syntenic units of the early diverging yeasts, like *trn*T-*trn*E-*trn*M1-*trn*M2, can be also found in mt genomes of species belonging to other subphyla such as Taphrinomycotina (this work) and all Pezizomycotina (Kouvelis et al., 2004; Pantou et al., 2008). This further indicates that these gene clusters were originally found in a common mitochondrial ancestor and became lost in more recently diverged taxa. At the same time, it designates the close relationship of these species with the Pezizomycotina subphylum. This conclusion agrees with previous phylogenetic studies which contributed to the examination of the synteny motifs in Pezizomycotina (Kouvelis et al., 2004; Pantou et al., 2008). If the above is combined with the results presented in this work, it becomes evident that small syntenic units, like *nad*4L-*nad*5, *atp*6-*atp*8 and *trn*S-*rnl*, are consistent ancestral elements, present in the mt genome of the fungal progenitor. Furthermore, this syntenic conservation may be additionally explained through a functional approach as was already shown in the case of metazoa (Taanman, 1999), where it was shown that small genes are co-translated in a polycistronic transcript. Therefore, the above synteny analysis leads to the conclusion that a confined “relaxed” gene order is retained throughout the yeast evolution.

The patterns of genome organization correlate with the tree topology. As previously mentioned, species that belong to group A present a different type of genome organization compared to species belonging to group B (Fig 3, Table S5, Table 2). The second group is composed of species that form the so-called “CUG-Ser1” clade and present conserved syntenic groups despite their major rearrangements. This high level of genetic rearrangements indicates that a significant amount of genetic shuffling took place during the alteration of the genetic code and formation of the CUG-Ser1 clade, at approximately 300 MYA (Fig 2, Table S5). This burst of genetic rearrangements is exclusively found in this clade. This is further supported by the fact that members of the Pichiaceae and CUG-Ala families (Groups C and D), which diverged from the CUG-Ser1 clade at a similar evolutionary time (325 MYA), present a partially different type of gene order and syntenic groups. Similarly, the major gene shuffling event that led to the formation of group A occurred during a similar time period (approximately 310 MYA) (Fig 2, Table S5). Still, the gene order of group A species was further differentiated with minimal late syntenic transpositions, leading to a variability which coincides chronologically firstly with the discrimination of Phaffomycetaceae from Saccharomycetaceae and secondly, to the split of Saccharomycetaceae into three clades (post-WGD, TZ and LKE). Therefore, the diverse synteny of mitogenomes found for taxa belonging to the same subgroups and at inter-species level, indicates that this genome shuffling is an ongoing process.

The rate of divergence in synteny and gene nucleotide composition (Fig 2) is not constant, but changes as the organisms evolve through time. This is detected if the time divergence of the families, examined from the mt dataset, is compared to the respective rate of the nuclear (nc) encoded dataset (Shen et al., 2018). Mt based analyses presented a slower rate of divergence compared to the respective values of nuclear datasets during the early evolution of yeasts, but in later evolutionary stages, the opposite was observed. For instance, the divergence of Pichiaceae and CUG-Ala clades from CUG-Ser1 (and

*H. uvarum*) was estimated at a time of 324 and 210 MYA based on the mt and nc datasets, respectively, while the split of the Saccharomycetaceae TZ clade from the post-WGD species were set to 75 and 93 MYA from the mt and nc datasets, respectively (Fig 2 and Shen et al., 2018). Therefore, in this work, a labile divergence rate of the mt genome evolution is presented and thus, conclusions proposed in other studies like the proposition that fungal mt genome evolution is slower to the respective nuclear genome (Sandor et al., 2018), must be approached cautiously.

The comparison of the mt tree (Fig 1) with the respective nuclear phylogeny, obtained for yeasts belonging to all known Saccharomycotina families (Shen et al., 2018), showed that both mt and nc datasets provide similar phylogenetic trees with excellent statistical support. Although both phylogenies are crucial for the unraveling of Saccharomycotina evolution, mitogenome phylogenetic analysis is faster and equally reliable compared to the nuclear one, as it resulted by the concatenation of only 14 mt proteins, whereas the nc dataset included 2,408 amino acid orthologous groups (Shen et al., 2018). Thus, it becomes evident that mitogenome based phylogenies may be highly informative for providing accurate and easily defined phylogenies.

Yet, throughout the evolution of yeasts, the mt genome content seems to be consistent in respect to its gene content, with the exception of the NADH subunit genes and *rps*3 (Table S1). The latter is a gene which was shown to be highly mobile in the kingdom of Fungi (Korovesi et al., 2018). In yeasts, there are only two groups, C (Pichiacheae) and D (CUG-Ser1 clade), in which the *rps*3 gene is absent, a fact possibly related to the genetic code alteration (Korovesi et al., 2018). According to previous studies that tried to explain the codon reassignments in mt genetic codes of eukaryotes, several mechanisms have been suggested to explain these alterations (Sengupta et al., 2005; Sengupta et al., 2007; Sengupta and Higgs, 2015). In detail, four mechanisms, i.e., “Codon disappearance”, the “unassigned codon”, the “ambiguous intermediate” and the “compensatory change”, have been proposed earlier (Sengupta et al., 2007), with the first two recently receiving more acceptance (Sengupta and Higgs, 2015; Koonin and Novozhilo, 2009). In our study, it became evident that the “Codon disappearance” mechanism might have driven evolution towards the observed genetic code changes through mutation and selection, as several codons, like ATA, CTN and TGG, are underrepresented as shown by the RCSU index in this work (Table S3) and another study (LaBella et al., 2019).

A comparison with the phylogenetic trees produced by mt phylogeny is needed to fully interpret genetic code alterations. Results from this study are in accordance with previous data from Sengupta et al. (2007) that described the major mitochondrial genetic code alteration events that took place during the evolution of all main phylogenetic eukaryotic lineages. The molecular clock revealed by our work provides an additional opportunity to estimate the timeframe during which these changes occurred. In detail, the reassignment of UGA stop to the Trp codon occurred as two independent events in the Ascomycota lineage (Sengupta et al., 2007), but only once during Saccharomycotina evolution (Fig. 2). This occurred between 590 and 548 MYA, when the subphylum Saccharomycotina diverged from Taphrinomycotina (used as outgroup in this work). This codon reassignment also marked the deviation from the standard genetic code, since all yeasts examined, besides the ones belonging to the Taphrinomycotina subphylum, employ the alterative mt genetic codes 3 and 4 (Table S1). CTN reassignment Leu to Thr occurred as a unique event right before the emergence of the Saccharomycetaceae and Saccharomycodaceae clades which took place approximately 259 and 200 MYA and marked the important evolutionary event of genetic code alteration from genetic code 4 to 3 (Fig 2). This is further confirmed by comparing the high RSCU values of the respective CTN codons (genetic code 3) among all phylogenetic groups (Fig 4). CTN codon employment is more distinct in cases of CTT and CTA codons, since they present significantly higher RSCU values in the above-mentioned phylogenetic clades, compared with the majority of the remaining groups/families which employ the ACN codon for Threonine translation (Fig 4). Reassignments of ATA from Ile to Met occurred as two independent events throughout yeast evolution: a) the first event took place approximately 188 MYA within the LKE clade, right before the separation of the subclade consisting of *Eremothecium* spp. The rest of the members of the LKE clade were not subjected into that genetic code reassignment (Fig 2), b) the second event took place within the post-WGD clade of Saccharomycetes, after its separation from the TZ clade (Fig 2). Sengupta et al. (2007) placed this event in the beginning of post-WGD clade formation in order to exclude members of the LKE clade, but did not analyze data from any representative of the TZ clade, thus, no conclusions could be made regarding codon reassignment within this clade. Conclusions from our work are supported by the fact that nowadays, information on mitochondrial genomes of more species are available. Our codon frequency analysis showed a very low RSCU value of the ATA Met codon in the TZ clade, indicating that this genetic codon reassignment of Ile to Met did not occur in members of this clade. Thus, it may be assumed that this reassignment occurred at the beginning of post WGD clade formation in a relatively recent event, less than 75 MYA (Fig 2 and Fig 4).

## 5. Conclusions

In conclusion, the mt based phylogenetic analysis of Saccharomycotina yeasts representing 11 known families proved to be equally reliable, when compared to the respective nuclear genome-based analyses, with alignments of fewer genes needed. Moreover, the molecular time clock of the concatenated mt dataset showed that the evolution of yeasts is perpetual with different divergence rates, i.e., slower at the early stages of evolution and faster at later stages, when compared to nuclear based molecular dating. Mt genome divergence of yeasts follows the previously proposed “aenaeon” model which explains the intron mobility and its contribution to the mt genome diversity (Megarioti and Kouvelis, 2020), since intron abundance and intergenic region variability of mitogenomes proved to be labile and major contributors to the genetic diversity of yeasts. This diversity is further attributed to the major rearrangements caused by genetic recombinational events but are also constrained by the conservation of the few ancestral syntenic units found throughout the main yeast phylogenetic lineages. Finally, genetic code alterations do happen throughout the evolution among the different families of yeasts, even during recent events, like the codon reassignment of ATA from Ile to Met, which occurred before the post-WGD clade formation. Therefore, mitogenomic analyses are crucial in deciphering the evolution of yeasts.

## Appendix A. Supplementary material

The following are the supplementary data of this article:

Table S1

Table S2

Table S3

Table S4

Table S5

Table S6

File S1

File S2

Fig S1

## Supplementary Data

**Table S1: The species and strain identity used in this study.** The Accession Number of their mt genome, the family and Phylogenetic groups in which they are classified, the genetic code, the topology, size of mt genomes, the number, total and % size of introns, the total and % size of intergenic regions, the AT content, presence of *nad* genes and *rps*3 gene, the number of *trn*s and the number of GC repeats with 90% identity are further provided.

The distribution of taxa according to their mt genome size is shown at the far right end of this table.

Yes= exists, No=absent, sa=standing alone gene, C= circular, L=linear, N/A=Not Available

**Table S2: The re-annotated and *de novo* annotated species and strains, as well as their Accession number**.

**Table S3: Detailed representation of Met (ATG, ATA), Thr (CTN, ACN), Trp (TGG, TGA) codons and codon frequency (RSCU values) for mt genes *cob,cox*1-3 and *atp*6,8,9**. RSCU values were calculated separately in 141 taxa studied, for each codon/ gene. In addition, a mean RSCU value for each codon/ each whole genome is presented in a separate table, as well as mean RSCU values for codon/genetic code. The mean RSCU value of codons of genes examined from all species employing genetic codes 3 and 4 (separately) are provided at the far side of this sheet.

**Table S4: Timetree analysis using the RelTime method.** A timetree inferred using the Reltime method and the General Reversible Mitochondrial + Freq. model as conducted in MEGA7. The timetree was computed using 4 calibration constraints. The estimated log likelihood value is −291342.08. A discrete Gamma distribution was used to model evolutionary rate differences among sites [5 categories (+G, parameter = 0.6990)].

**Table S5: Detailed gene order (synteny) of species examined. Syntenic units are presented in different colors**. The pink color in species names represents species used for Figure 3.

**Table S6: I) Intron distribution among (a) families and phylogenetic groups/clades, (b) genes, (c) genes containing introns within Families or Phylogenetic clades, for the 141 taxa examined in this work**. Empty cells denote the absence of introns. **II) Gene, intron’s name, introns’ type, gene 5’ and 3’ insertion sites and sequence position, number of ORFs and ORF’s type for the phylogenetic clades: CUG-Ser2, Dipodascaceae/Trichomonascaceae, Saccharomycodaceae, CUG-Ala, Sporopachydermia, Lipomycetaceae and Trigonopsidaceae**. The intron analysis for the rest of the phylogenetic clades are already published in previous work (Megarioti and Kouvelis, 2020).

**Fig S1: Detailed time calibrated phylogeny of the Saccharomycotina subphylum.** The 141 mt genomes analyzed in this study are shown in color according to the families/ phylogenetic clades of the tree shown in Fig 1. In detail, Saccharomycetaceae blue, Saccharomycodaceae dark green, Phaffomycetaceae light blue, GUG-Ser1 clade yellow, CUG-Ser2 clade light green, Pichiaceae orange, Sporopachydermia clade dark purple, Dipodascaceae/ Trichomonascaceae clade red, Lipomycetaceae light purple, Trigonospidaceae pink, CUG-Ala clade brown and Taphrinomycotina black (outgroup). The time calibration bar is provided at the bottom of the figure.

**FiFile S1: NJ - single gene phylogenetic trees** based on each mitochondrial gene of all species examined in this study.

**File S2: Logos of the GC clusters for each representative species.**

## References

Abascal, F., Zardoya, R., & Posada, D. (2005). ProtTest: selection of best-fit models of protein evolution. Bioinformatics (Oxford, England), 21(9), 2104–2105. https://doi.org/10.1093/bioinformatics/bti263.

Altschul, S. F., Gish, W., Miller, W., Myers, E. W., & Lipman, D. J. (1990). Basic local alignment search tool. Journal of molecular biology, 215(3), 403–410. https://doi.org/10.1016/S0022-2836(05)80360-2

Ballard JWO, and Pichaud N (2014). Mitochondrial DNA: More than an evolutionary bystander. Functional Ecology 28:218–231. doi: 10.1111/1365-2435.12177

Ballard, J. W., & Whitlock, M. C. (2004). The incomplete natural history of mitochondria. Molecular ecology, 13(4), 729–744. https://doi.org/10.1046/j.1365-294x.2003.02063.x.

Barreto, F. S., Watson, E. T., Lima, T. G., Willett, C. S., Edmands, S., Li, W., & Burton, R. S. (2018). Genomic signatures of mitonuclear coevolution across populations of *Tigriopus californicus*. Nature ecology & evolution, 2(8), 1250–1257. https://doi.org/10.1038/s41559-018-0588-1

Bartelli, T. F., Ferreira, R. C., Colombo, A. L., & Briones, M. R. (2013). Intraspecific comparative genomics of *Candida albicans* mitochondria reveals non-coding regions under neutral evolution. Infection, genetics and evolution: journal of molecular epidemiology and evolutionary genetics in infectious diseases, 14, 302–312. https://doi.org/10.1016/j.meegid.2012.12.012

Benítez, T., Martínez, P., & Codón, A. C. (1996). Genetic constitution of industrial yeast. Microbiologia (Madrid, Spain), 12(3), 371–384.

Botstein, D., & Fink, G. R. (2011). Yeast: an experimental organism for 21st Century biology. Genetics, 189(3), 695–704. https://doi.org/10.1534/genetics.111.130765

Bouchier, C., Ma, L., Créno, S., Dujon, B., & Fairhead, C. (2009). Complete mitochondrial genome sequences of three *Nakaseomyces* species reveal invasion by palindromic GC clusters and considerable size expansion. FEMS yeast research, 9(8), 1283–1292. https://doi.org/10.1111/j.1567-1364.2009.00551.x

Brankovics, B., van Dam, P., Rep, M., de Hoog, G. S., J van der Lee, T. A., Waalwijk, C., & van Diepeningen, A. D. (2017). Mitochondrial genomes reveal recombination in the presumed asexual *Fusarium oxysporum* species complex. BMC genomics, 18(1), 735. https://doi.org/10.1186/s12864-017-4116-5

Braun E. L. (2003). Innovation from reduction: gene loss, domain loss and sequence divergence in genome evolution. Applied bioinformatics, 2(1), 13–34.

Burger, G., Gray, M. W., & Lang, B. F. (2003). Mitochondrial genomes: anything goes. Trends in genetics: TIG, 19(12), 709–716. https://doi.org/10.1016/j.tig.2003.10.012

Burland T. G. (2000). DNASTAR’s Lasergene sequence analysis software. Methods in molecular biology (Clifton, N.J.), 132, 71–91. https://doi.org/10.1385/1-59259-192-2:71

Chan, P. P., & Lowe, T. M. (2019). tRNAscan-SE: Searching for tRNA Genes in Genomic Sequences. Methods in molecular biology (Clifton, N.J.), 1962, 1–14. https://doi.org/10.1007/978-1-4939-9173-0_1

Crooks, G. E., Hon, G., Chandonia, J. M., & Brenner, S. E. (2004). WebLogo: a sequence logo generator. Genome research, 14(6), 1188–1190. https://doi.org/10.1101/gr.849004

De Chiara, M., Friedrich, A., Barré, B., Breitenbach, M., Schacherer, J., & Liti, G. (2020). Discordant evolution of mitochondrial and nuclear yeast genomes at population level. BMC biology, 18(1), 49. https://doi.org/10.1186/s12915-020-00786-4

de Zamaroczy, M., & Bernardi, G. (1985). Sequence organization of the mitochondrial genome of yeast-a review. Gene, 37(1-3), 1–17. https://doi.org/10.1016/0378-1119(85)90252-5

de Zamaroczy, M., & Bernardi, G. (1986). The GC clusters of the mitochondrial genome of yeast and their evolutionary origin. Gene, 41(1), 1–22. https://doi.org/10.1016/0378-1119(86)90262-3

Deng, Y., Hsiang, T., Li, S., Lin, L., Wang, Q., Chen, Q., Xie, B., & Ming, R. (2018). Comparison of the Mitochondrial Genome Sequences of Six *Annulohypoxylon stygium* Isolates Suggests Short Fragment Insertions as a Potential Factor Leading to Larger Genomic Size. Frontiers in microbiology, 9, 2079. https://doi.org/10.3389/fmicb.2018.02079

Dieckmann, C. L., & Gandy, B. (1987). Preferential recombination between GC clusters in yeast mitochondrial DNA. The EMBO journal, 6(13), 4197–4203.

Dujon B. (2006). Yeasts illustrate the molecular mechanisms of eukaryotic genome evolution. Trends in genetics: TIG, 22(7), 375–387. https://doi.org/10.1016/j.tig.2006.05.007

Dujon B. (2010). Yeast evolutionary genomics. Nature reviews. Genetics, 11(7), 512–524. https://doi.org/10.1038/nrg2811

Duò, A., Bruggmann, R., Zoller, S., Bernt, M., & Grünig, C. R. (2012). Mitochondrial genome evolution in species belonging to the *Phialocephala fortinii* s.l. - *Acephala applanata* species complex. BMC genomics, 13, 166. https://doi.org/10.1186/1471-2164-13-166

Eldarov, M. A., Kishkovskaia, S. A., Tanaschuk, T. N., & Mardanov, A. V. (2016). Genomics and Biochemistry of *Saccharomyces cerevisiae* Wine Yeast Strains. Biochemistry. Biokhimiia, 81(13), 1650–1668. https://doi.org/10.1134/S0006297916130046

Escalera-Fanjul X., Quezada, H., Riego-Ruiz L., & González, A. (2019). Whole-Genome Duplication and Yeast’s Fruitful Way of Life. Trends in genetics: TIG, 35(1), 42–54. https://doi.org/10.1016/j.tig.2018.09.008

Fisher, K. J., Buskirk, S. W., Vignogna, R. C., Marad, D. A., & Lang, G. I. (2018). Adaptive genome duplication affects patterns of molecular evolution in *Saccharomyces cerevisiae*. PLoS genetics, 14(5), e1007396. https://doi.org/10.1371/journal.pgen.1007396

Foury, F., Roganti, T., Lecrenier, N., & Purnelle, B. (1998). The complete sequence of the mitochondrial genome of *Saccharomyces cerevisiae*. FEBS letters, 440(3), 325–331. https://doi.org/10.1016/s0014-5793(98)01467-7

Freel, K. C., Friedrich, A., & Schacherer, J. (2015). Mitochondrial genome evolution in yeasts: an all-encompassing view. FEMS yeast research, 15(4), fov023. https://doi.org/10.1093/femsyr/fov023

Fricova, D., Valach, M., Farkas, Z., Pfeiffer, I., Kucsera, J., Tomaska, L., & Nosek, J. (2010). The mitochondrial genome of the pathogenic yeast *Candida subhashii*: GC-rich linear DNA with a protein covalently attached to the 5′ termini. Microbiology (Reading, England), 156(7), 2153–2163. https://doi.org/10.1099/mic.0.038646-0

Fritsch, E. S., Chabbert, C. D., Klaus, B., & Steinmetz, L. M. (2014). A genome-wide map of mitochondrial DNA recombination in yeast. Genetics, 198(2), 755–771. https://doi.org/10.1534/genetics.114.166637

Hatefi Y. (1985). The mitochondrial electron transport and oxidative phosphorylation system. Annual review of biochemistry, 54, 1015–1069. https://doi.org/10.1146/annurev.bi.54.070185.005055

Hill G. E. (2020). Genetic hitchhiking, mitonuclear coadaptation, and the origins of mt DNA barcode gaps. Ecology and evolution, 10(17), 9048–9059. https://doi.org/10.1002/ece3.6640

Huang, Y., Niu, B., Gao, Y., Fu, L., & Li, W. (2010). CD-HIT Suite: a web server for clustering and comparing biological sequences. Bioinformatics (Oxford, England), 26(5), 680–682. https://doi.org/10.1093/bioinformatics/btq003

Kellis, M., Birren, B. W., & Lander, E. S. (2004). Proof and evolutionary analysis of ancient genome duplication in the yeast *Saccharomyces cerevisiae*. Nature, 428(6983), 617–624. https://doi.org/10.1038/nature02424

Knop M. (2011). Yeast cell morphology and sexual reproduction-a short overview and some considerations. Comptes rendus biologies, 334(8-9), 599–606. https://doi.org/10.1016/j.crvi.2011.05.007

Koonin, E. V., & Novozhilov, A. S. (2009). Origin and evolution of the genetic code: the universal enigma. IUBMB life, 61(2), 99–111. https://doi.org/10.1002/iub.146

Korovesi, A. G., Ntertilis, M., & Kouvelis, V. N. (2018). Mt-*rps*3 is an ancient gene which provides insight into the evolution of fungal mitochondrial genomes. Molecular phylogenetics and evolution, 127, 74–86. https://doi.org/10.1016/j.ympev.2018.04.037

Kortsinoglou, A. M., Korovesi, A. G., Theelen, B., Hagen, F., Boekhout, T., & Kouvelis, V. N. (2019). The mitochondrial intergenic regions *nad*1-*cob* and *cob*-*rps*3 as molecular identification tools for pathogenic members of the genus *Cryptococcus*. FEMS yeast research, 19(8), foz077. https://doi.org/10.1093/femsyr/foz077

Kouvelis, V. N., Ghikas, D. V., & Typas, M. A. (2004). The analysis of the complete mitochondrial genome of *Lecanicillium muscarium* (synonym *Verticillium lecanii*) suggests a minimum common gene organization in mtDNAs of Sordariomycetes: phylogenetic implications. Fungal genetics and biology: FG & B, 41(10), 930–940. https://doi.org/10.1016/j.fgb.2004.07.003

Kouvelis, V. N., Sialakouma, A., & Typas, M. A. (2008). Mitochondrial gene sequences alone or combined with ITS region sequences provide firm molecular criteria for the classification of Lecanicillium species. Mycological research, 112(Pt 7), 829–844. https://doi.org/10.1016/j.mycres.2008.01.016

Kovác, L., Lazowska, J., & Slonimski, P. P. (1984). A yeast with linear molecules of mitochondrial DNA. Molecular & general genetics: MGG, 197(3), 420–424. https://doi.org/10.1007/BF00329938

Krassowski, T., Coughlan, A. Y., Shen, X. X., Zhou, X., Kominek, J., Opulente, D. A., Riley, R., Grigoriev, I. V., Maheshwari, N., Shields, D. C., Kurtzman, C. P., Hittinger, C. T., Rokas, A., & Wolfe, K. H. (2018). Evolutionary instability of CUG-Leu in the genetic code of budding yeasts. Nature communications, 9(1), 1887. https://doi.org/10.1038/s41467-018-04374-7

Kumar, S., Stecher, G., & Tamura, K. (2016). MEGA7: Molecular Evolutionary Genetics Analysis Version 7.0 for Bigger Datasets. Molecular biology and evolution, 33(7), 1870–1874. https://doi.org/10.1093/molbev/msw054

Kumar, S., Stecher, G., Suleski, M., & Hedges, S. B. (2017). TimeTree: A Resource for Timelines, Timetrees, and Divergence Times. Molecular biology and evolution, 34(7), 1812–1819. https://doi.org/10.1093/molbev/msx116

Kurtzman C. P. (2011). Phylogeny of the ascomycetous yeasts and the renaming of *Pichia anomala* to *Wickerhamomyces anomalus*. Antonie van Leeuwenhoek, 99(1), 13–23. https://doi.org/10.1007/s10482-010-9505-6

Kurtzman, C. P., & Boekhout, T. (2017). Yeasts as Distinct Life Forms of Fungi. In: Buzzini, P., Lachance, M. A., Yurkov, A. (eds) Yeasts in Natural Ecosystems: Ecology. Springer, Cham. https://doi.org/10.1007/978-3-319-61575-2_1

LaBella, A. L., Opulente, D. A., Steenwyk, J. L., Hittinger, C. T., & Rokas, A. (2019). Variation and selection on codon usage bias across an entire subphylum. PLoS genetics, 15(7), e1008304. https://doi.org/10.1371/journal.pgen.1008304

Lang, B. F., Gray, M. W., & Burger, G. (1999). Mitochondrial genome evolution and the origin of eukaryotes. Annual review of genetics, 33, 351–397. https://doi.org/10.1146/annurev.genet.33.1.351

Lang, B. F., Laforest, M. J., & Burger, G. (2007). Mitochondrial introns: a critical view. Trends in genetics: TIG, 23(3), 119–125. https://doi.org/10.1016/j.tig.2007.01.006

Madeira, F., Park, Y. M., Lee, J., Buso, N., Gur, T., Madhusoodanan, N., Basutkar, P., Tivey, A., Potter, S. C., Finn, R. D., & Lopez, R. (2019). The EMBL-EBI search and sequence analysis tools APIs in 2019. Nucleic acids research, 47(W1), W636–W641. https://doi.org/10.1093/nar/gkz268

Marcet-Houben M., & Gabaldón, T. (2015). Beyond the Whole-Genome Duplication: Phylogenetic Evidence for an Ancient Interspecies Hybridization in the Baker’s Yeast Lineage. PLoS biology, 13(8), e1002220. https://doi.org/10.1371/journal.pbio.1002220

Masel J. (2011). Genetic drift. Current biology: CB, 21(20), R837–R838. https://doi.org/10.1016/j.cub.2011.08.007

Megarioti, A. H., & Kouvelis, V. N. (2020). The Coevolution of Fungal Mitochondrial Introns and Their Homing Endonucleases (GIY-YIG and LAGLIDADG). Genome biology and evolution, 12(8), 1337–1354. https://doi.org/10.1093/gbe/evaa126

Meiklejohn, C. D., Montooth, K. L., & Rand, D. M. (2007). Positive and negative selection on the mitochondrial genome. Trends in genetics: TIG, 23(6), 259–263. https://doi.org/10.1016/j.tig.2007.03.008

Miranda, I., Silva, R., & Santos, M. A. (2006). Evolution of the genetic code in yeasts. Yeast (Chichester, England), 23(3), 203–213. https://doi.org/10.1002/yea.1350

Nielsen J. (2019). Yeast Systems Biology: Model Organism and Cell Factory. Biotechnology journal, 14(9), e1800421. https://doi.org/10.1002/biot.201800421

Nosek, J., & Tomáska, L. (2003). Mitochondrial genome diversity: evolution of the molecular architecture and replication strategy. Current genetics, 44(2), 73–84. https://doi.org/10.1007/s00294-003-0426-z

Nosek, J., Tomáska, L., Fukuhara, H., Suyama, Y., & Kovác, L. (1998). Linear mitochondrial genomes: 30 years down the line. Trends in genetics: TIG, 14(5), 184–188. https://doi.org/10.1016/s0168-9525(98)01443-7

Okonechnikov, K., Golosova, O., Fursov, M., & UGENE team (2012). Unipro UGENE: a unified bioinformatics toolkit. Bioinformatics (Oxford, England), 28(8), 1166–1167. https://doi.org/10.1093/bioinformatics/bts091

Pantou, M. P., Kouvelis, V. N., & Typas, M. A. (2008). The complete mitochondrial genome of *Fusarium oxysporum*: insights into fungal mitochondrial evolution. Gene, 419(1-2), 7–15. https://doi.org/10.1016/j.gene.2008.04.009

Peter, J., De Chiara, M., Friedrich, A., Yue, J. X., Pflieger, D., Bergström, A., Sigwalt, A., Barre, B., Freel, K., Llored, A., Cruaud, C., Labadie, K., Aury, J. M., Istace, B., Lebrigand, K., Barbry, P., Engelen, S., Lemainque, A., Wincker, P., Liti, G., … Schacherer, J. (2018). Genome evolution across 1,011 *Saccharomyces cerevisiae* isolates. Nature, 556(7701), 339–344. https://doi.org/10.1038/s41586-018-0030-5

Pote, S. T., Sonawane, M. S., Rahi, P., Shah, S. R., Shouche, Y. S., Patole, M. S., Thakar, M. R., & Sharma, R. (2020). Distribution of Pathogenic Yeasts in Different Clinical Samples: Their Identification, Antifungal Susceptibility Pattern, and Cell Invasion Assays. Infection and drug resistance, 13, 1133–1145. https://doi.org/10.2147/IDR.S238002

Pramateftaki, P. V., Kouvelis, V. N., Lanaridis, P., & Typas, M. A. (2006). The mitochondrial genome of the wine yeast *Hanseniaspora uvarum*: a unique genome organization among yeast/fungal counterparts. FEMS yeast research, 6(1), 77–90. https://doi.org/10.1111/j.1567-1364.2005.00018.x

Ronquist, F., & Huelsenbeck, J. P. (2003). MrBayes 3: Bayesian phylogenetic inference under mixed models. Bioinformatics (Oxford, England), 19(12), 1572–1574. https://doi.org/10.1093/bioinformatics/btg180

Saccone, C., De Giorgi, C., Gissi, C., Pesole, G., & Reyes, A. (1999). Evolutionary genomics in Metazoa: the mitochondrial DNA as a model system. Gene, 238(1), 195–209. https://doi.org/10.1016/s0378-1119(99)00270-x

Sandor, S., Zhang, Y., & Xu, J. (2018). Fungal mitochondrial genomes and genetic polymorphisms. Applied microbiology and biotechnology, 102(22), 9433–9448. https://doi.org/10.1007/s00253-018-9350-5

Santos, M. A., Ueda, T., Watanabe, K., & Tuite, M. F. (1997). The non-standard genetic code of *Candida* spp.: an evolving genetic code or a novel mechanism for adaptation?. Molecular microbiology, 26(3), 423–431. https://doi.org/10.1046/j.1365-2958.1997.5891961.x

Sekito, T., Okamoto, K., Kitano, H., & Yoshida, K. (1995). The complete mitochondrial DNA sequence of *Hansenula wingei* reveals new characteristics of yeast mitochondria. Current genetics, 28(1), 39–53. https://doi.org/10.1007/BF00311880

Sengupta, S., & Higgs, P. G. (2005). A unified model of codon reassignment in alternative genetic codes. Genetics, 170(2), 831–840. https://doi.org/10.1534/genetics.104.037887

Sengupta, S., & Higgs, P. G. (2015). Pathways of Genetic Code Evolution in Ancient and Modern Organisms. Journal of molecular evolution, 80(5-6), 229–243. https://doi.org/10.1007/s00239-015-9686-8

Sengupta, S., Yang, X., & Higgs, P. G. (2007). The mechanisms of codon reassignments in mitochondrial genetic codes. Journal of molecular evolution, 64(6), 662–688. https://doi.org/10.1007/s00239-006-0284-7

Sharp, P. M., Tuohy, T. M., & Mosurski, K. R. (1986). Codon usage in yeast: cluster analysis clearly differentiates highly and lowly expressed genes. Nucleic acids research, 14(13), 5125–5143. https://doi.org/10.1093/nar/14.13.5125

Shen, X. X., Opulente, D. A., Kominek, J., Zhou, X., Steenwyk, J. L., Buh, K. V., Haase, M., Wisecaver, J. H., Wang, M., Doering, D. T., Boudouris, J. T., Schneider, R. M., Langdon, Q. K., Ohkuma, M., Endoh, R., Takashima, M., Manabe, R. I., Čadež, N., Libkind, D., Rosa, C. A., … Rokas, A. (2018). Tempo and Mode of Genome Evolution in the Budding Yeast Subphylum. Cell, 175(6), 1533–1545.e20. https://doi.org/10.1016/j.cell.2018.10.023

Smith, J. M., & Haigh, J. (1974). The hitch-hiking effect of a favourable gene. Genetical research, 23(1), 23–35.

Solieri L. (2010). Mitochondrial inheritance in budding yeasts: towards an integrated understanding. Trends in microbiology, 18(11), 521–530. https://doi.org/10.1016/j.tim.2010.08.001

Stajich, J. E., Berbee, M. L., Blackwell, M., Hibbett, D. S., James, T. Y., Spatafora, J. W., & Taylor, J. W. (2009). The fungi. Current biology: CB, 19(18), R840–R845. https://doi.org/10.1016/j.cub.2009.07.004

Stamatakis A. (2014). RAxML version 8: a tool for phylogenetic analysis and post-analysis of large phylogenies. Bioinformatics (Oxford, England), 30(9), 1312–1313. https://doi.org/10.1093/bioinformatics/btu033

Stein, A., & Sia, E. A. (2017). Mitochondrial DNA repair and damage tolerance. Frontiers in bioscience (Landmark edition), 22, 920–943. https://doi.org/10.2741/4525

Stothard P. (2000). The sequence manipulation suite: JavaScript programs for analyzing and formatting protein and DNA sequences. BioTechniques, 28(6), 1102–1104. https://doi.org/10.2144/00286ir01

Taanman J. W. (1999). The mitochondrial genome: structure, transcription, translation and replication. Biochimica et biophysica acta, 1410(2), 103–123. https://doi.org/10.1016/s0005-2728(98)00161-3

Teresa Fernández-Espinar, M., Barrio, E., & Querol, A. (2003). Analysis of the genetic variability in the species of the *Saccharomyces* sensu stricto complex. Yeast (Chichester, England), 20(14), 1213–1226. https://doi.org/10.1002/yea.1034

Thompson, J. D., Higgins, D. G., & Gibson, T. J. (1994). CLUSTAL W: improving the sensitivity of progressive multiple sequence alignment through sequence weighting, position-specific gap penalties and weight matrix choice. Nucleic acids research, 22(22), 4673–4680. https://doi.org/10.1093/nar/22.22.4673

Wang, J. M., Bennett, R. J., & Anderson, M. Z. (2018). The Genome of the Human Pathogen *Candida albicans* Is Shaped by Mutation and Cryptic Sexual Recombination. mBio, 9(5), e01205-18. https://doi.org/10.1128/mBio.01205-18

Wesolowski, M., & Fukuhara, H. (1981). Linear mitochondrial deoxyribonucleic acid from the yeast *Hansenula mrakii*. Molecular and cellular biology, 1(5), 387–393. https://doi.org/10.1128/mcb.1.5.387-393.1981

Wilgenbusch, J. C., & Swofford, D. (2003). Inferring evolutionary trees with PAUP*. Current protocols in bioinformatics, Chapter 6. https://doi.org/10.1002/0471250953.bi0604s00

Williamson D. (2002). The curious history of yeast mitochondrial DNA. Nature reviews. Genetics, 3(6), 475–481. https://doi.org/10.1038/nrg814

Wolfe K. H. (2015). Origin of the Yeast Whole-Genome Duplication. PLoS biology, 13(8), e1002221. https://doi.org/10.1371/journal.pbio.1002221

Wolfe, K. H., & Shields, D. C. (1997). Molecular evidence for an ancient duplication of the entire yeast genome. Nature, 387(6634), 708–713. https://doi.org/10.1038/42711

Wolters, J. F., Chiu, K., & Fiumera, H. L. (2015). Population structure of mitochondrial genomes in Saccharomyces cerevisiae. BMC genomics, 16(1), 451. https://doi.org/10.1186/s12864-015-1664-4

Xiao, S., Nguyen, D. T., Wu, B., & Hao, W. (2017). Genetic Drift and Indel Mutation in the Evolution of Yeast Mitochondrial Genome Size. Genome biology and evolution, 9(11), 3088–3099. https://doi.org/10.1093/gbe/evx232

