## Supplementary figures and images for "Mitogenomics and mitochondrial gene phylogeny decipher the evolution of Saccharomycotina yeasts"

### Supplemental Figure S1

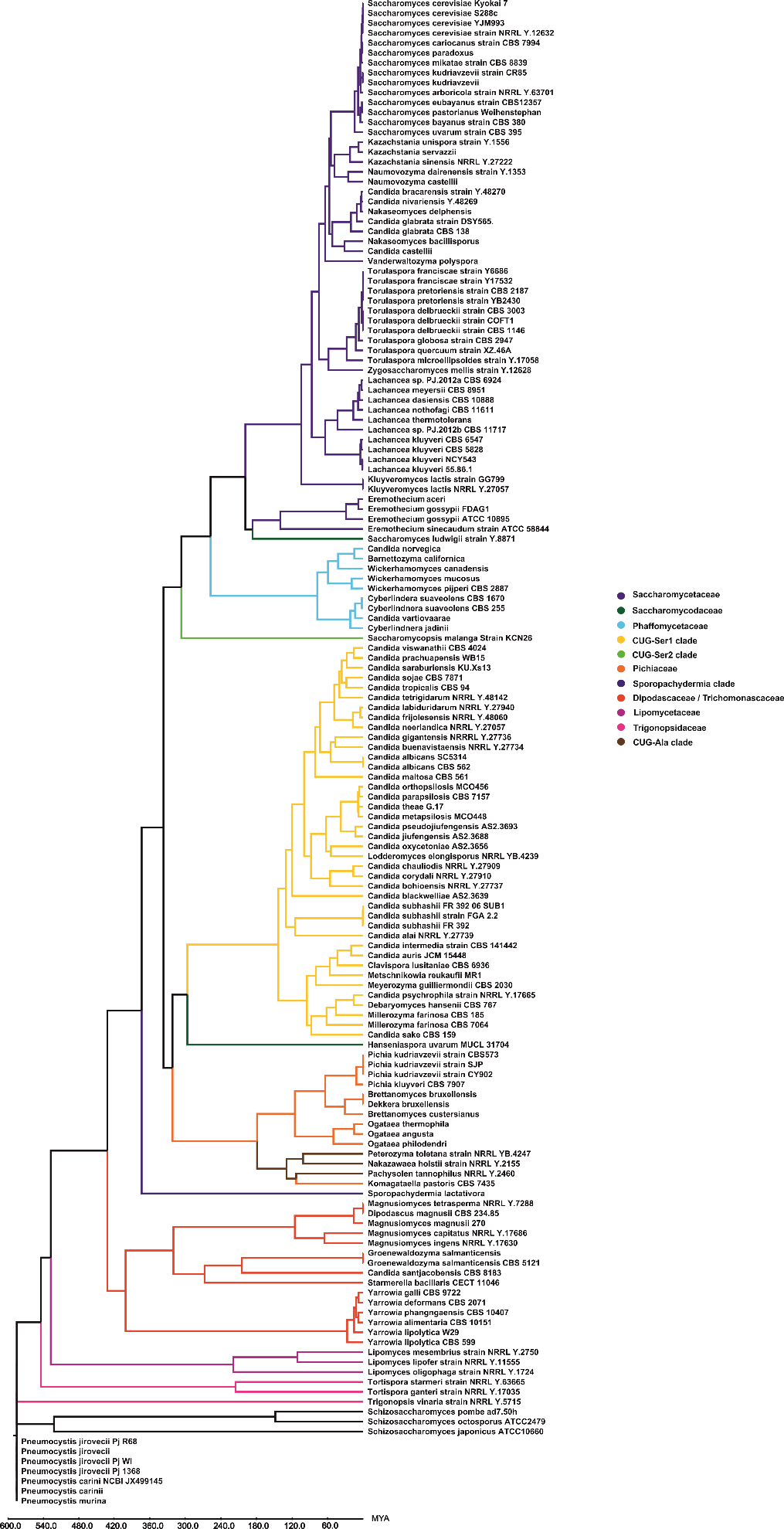
