## Supplemental File S2 for "Mitogenomics and mitochondrial gene phylogeny decipher the evolution of Saccharomycotina yeasts"

**S2 File: Logos of the GC clusters for each representative species.**

### Dipodascaceae/Trichomoscaceae clade

*Yarrowia prachuapensis* (435 repeats)

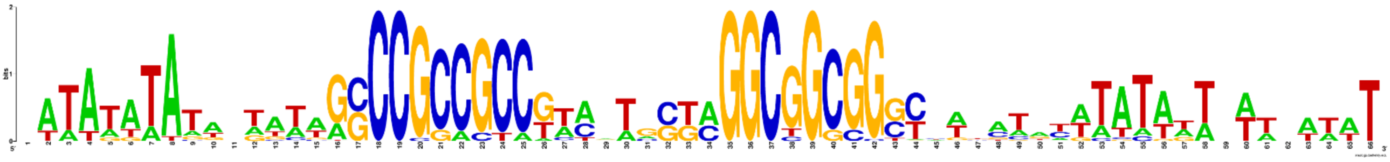

*Magnusiomyces tetrasperma* (11 repeats)

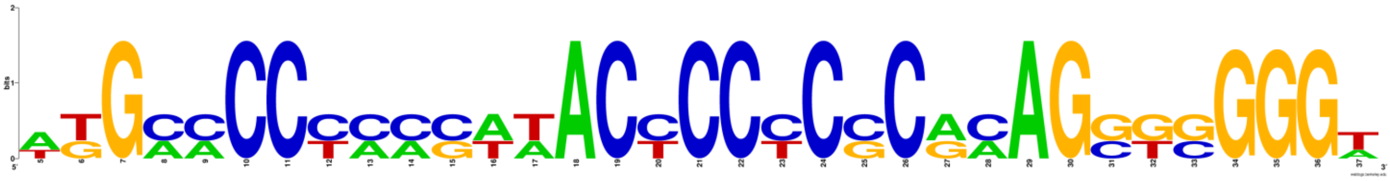

*Cyberlindnera suaveolens* strain CBS 1670 (2 repeats)

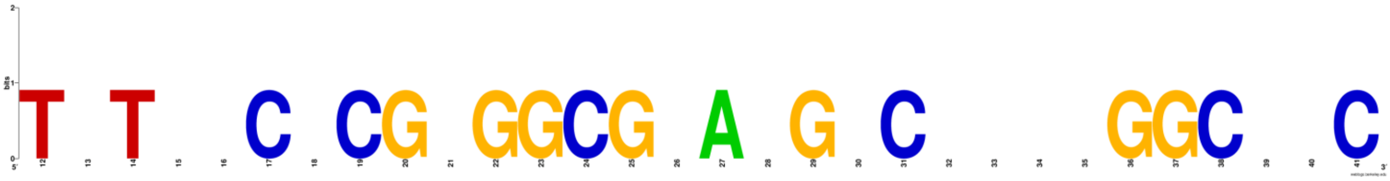

### CUG-Ser1 clade

*Candida gigantensis* strain NRRL Y-27736 (98 repeats)

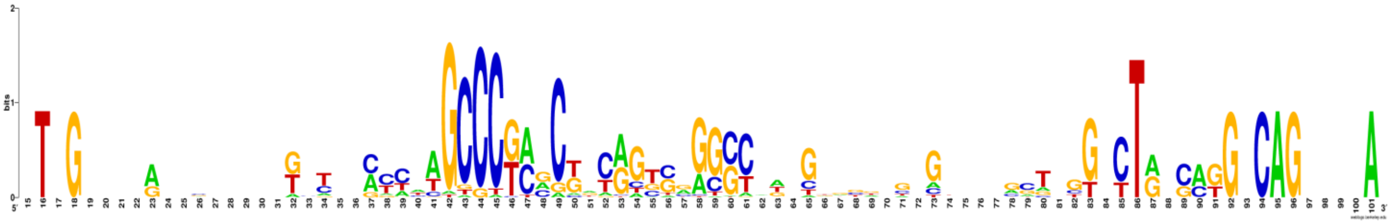

*Candida buenavistaensis* strain NRRL Y-27734 (328 repeats)

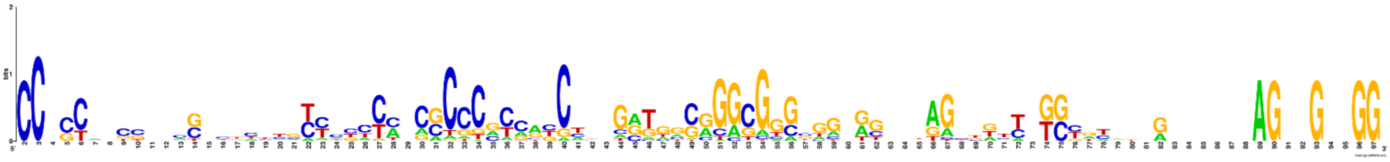

*Candida chauliodes* strain NRRL Y-27909 (245 repeats)

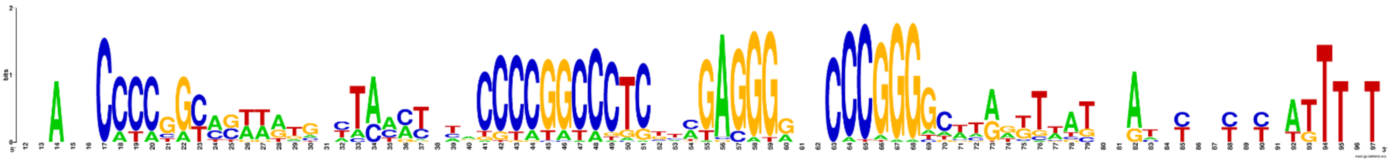

*Candida tropicalis* strain CBS94 (322 repeats)

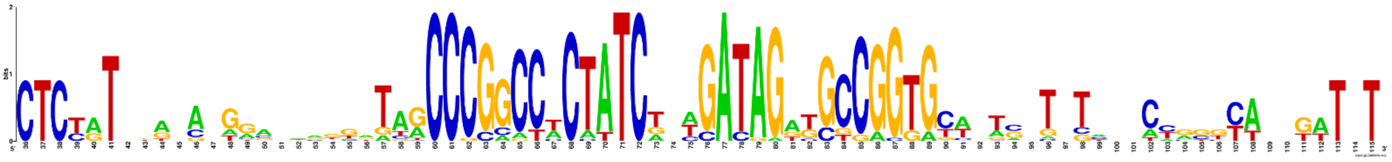

*Candida bohiensis* (339 repeats)

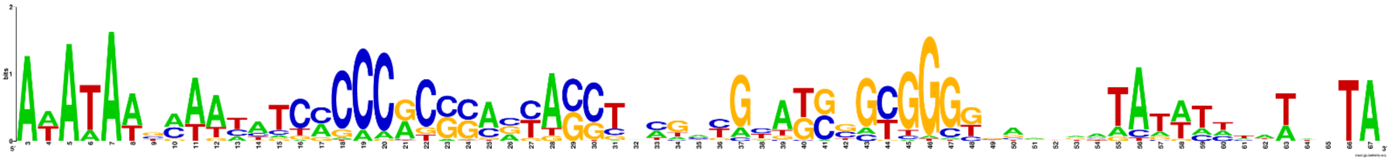

*Candida maltose* (396 repeats)

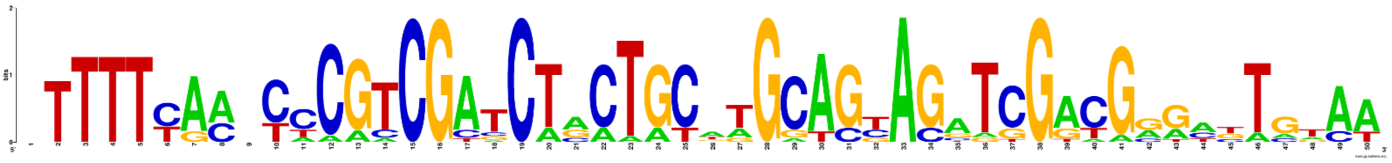

*Candida oxycetoniae* strain AS2.3656 (25 repeats)

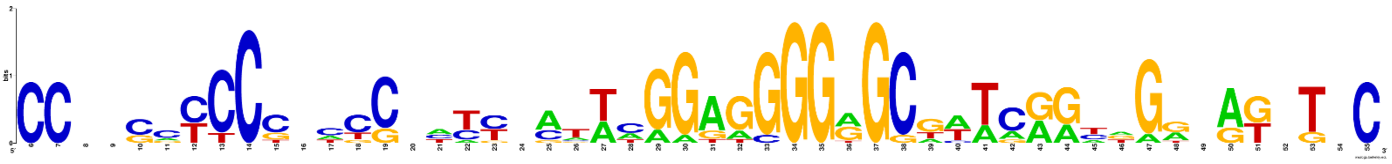

*Candida albicans* strain L757 (16 repeats)

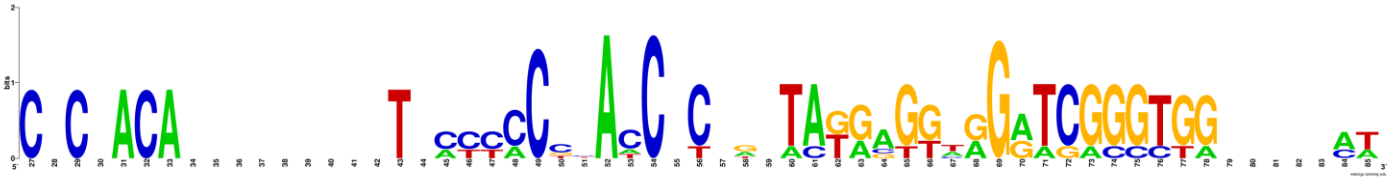

*Candida blackwellae* strain AS2.3639 (66 repeats)

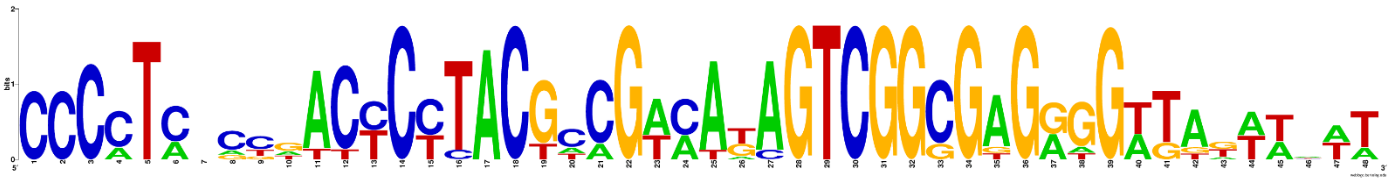

*Candida albicans* SC5314 (16 repeats)

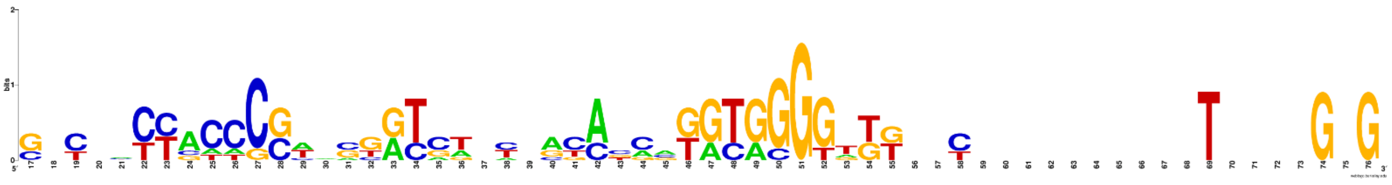

*Candida subhashii* strain FGA (8 repeats)

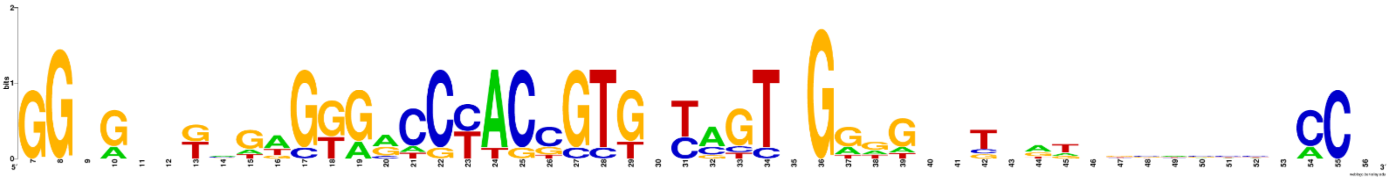

*Candida subhashii* strain FR-392-06-SUB1 (7 repeats)

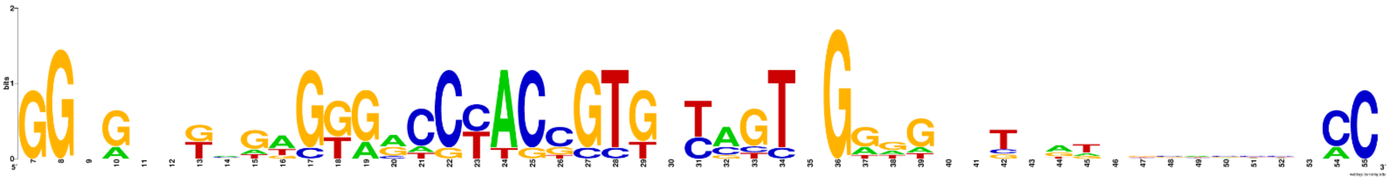

*Candida subhashii* (8 repeats)

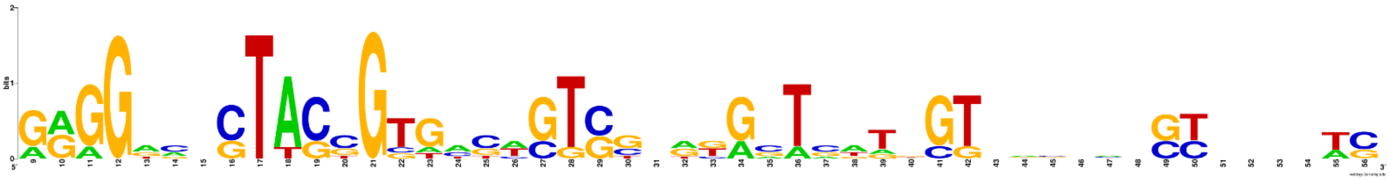

*Metschnikowia reukaufii* strain MR1 (7 repeats)

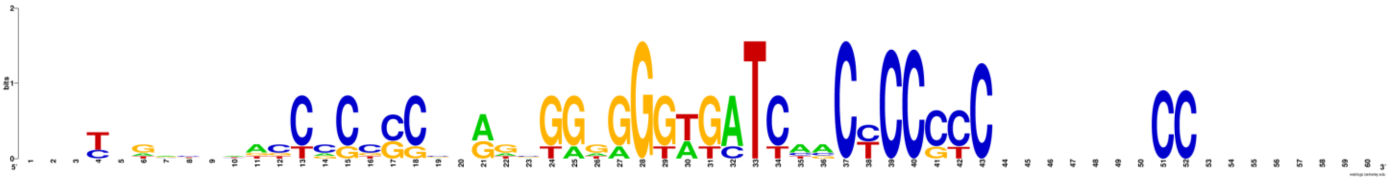

*Candida sojae* (12 repeats)

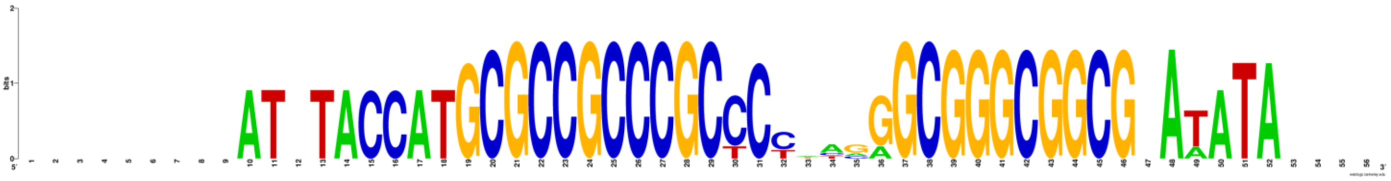

*Candida sake* strain CBS 15 (6 repeats)

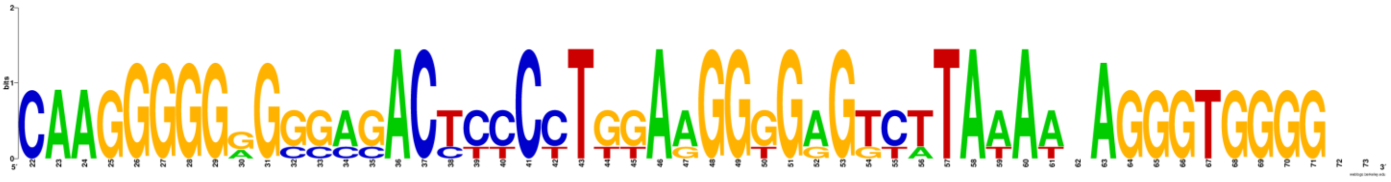

*Candida castellii* (9 repeats)

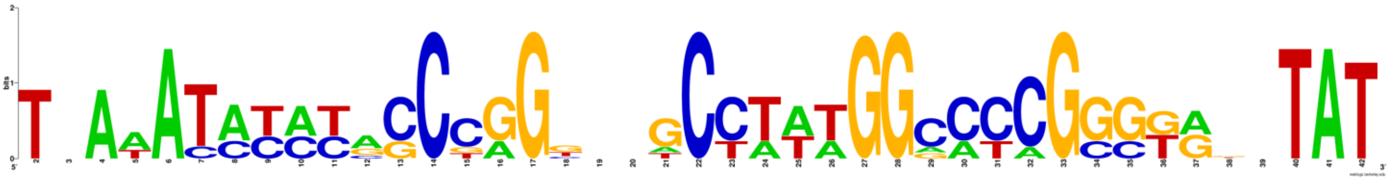

*Candida saraburiensis* strain KU-Xs13 (5 repeats)

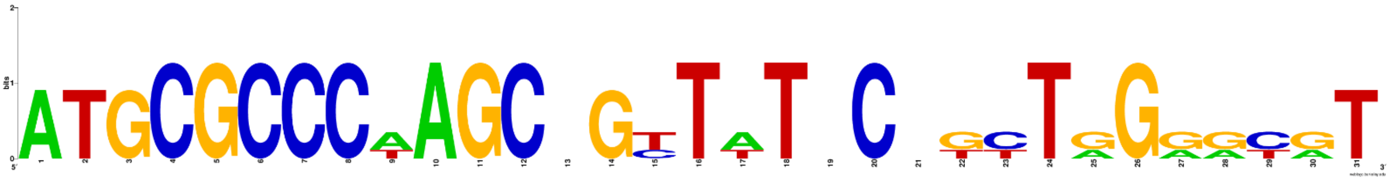

### Saccharomycetaceae post-WGD clade

*Saccharomyces pastorianus* Weihenstephan 34 70 (56 repeats)

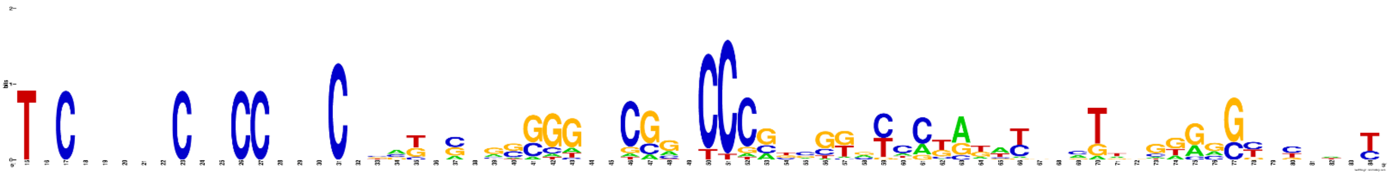

*Saccharomyces cerevisiae* YJM993 (194 repeats)

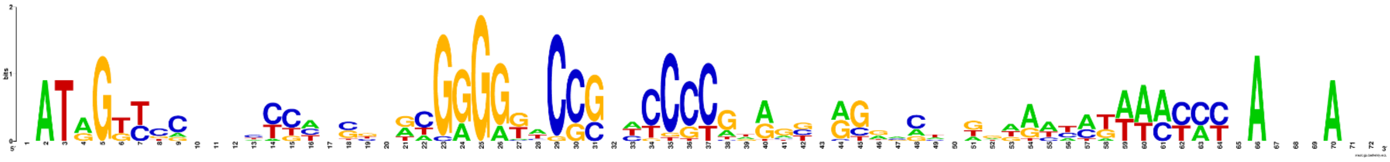

*Saccharomyces cerevisiae* strain NRRL Y-12632 (234 repeats)

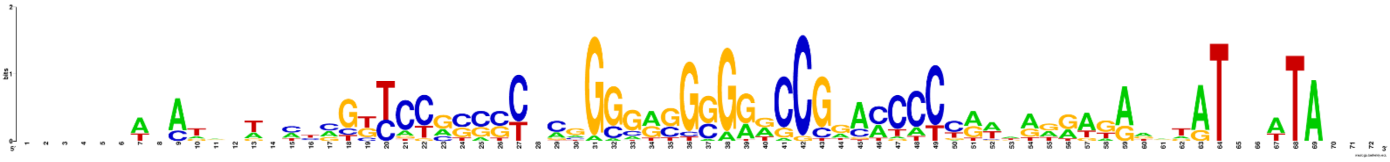

*Saccharomyces cerevisiae* S288c (228 repeats)

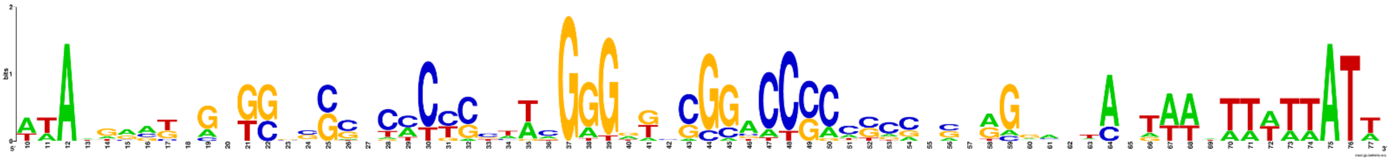

*Saccharomyces cerevisiae* Kyokai no. 7 (175 repeats)

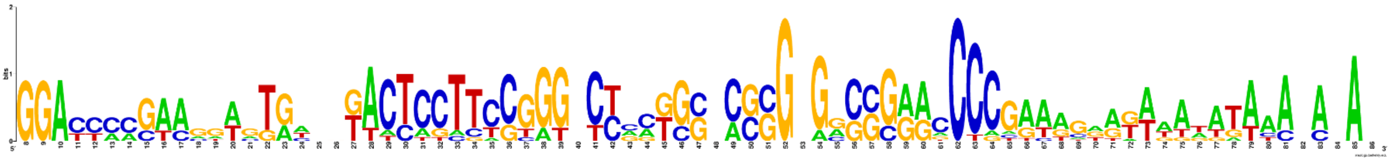

*Saccharomyces kudriavzevii* (60 repeats)

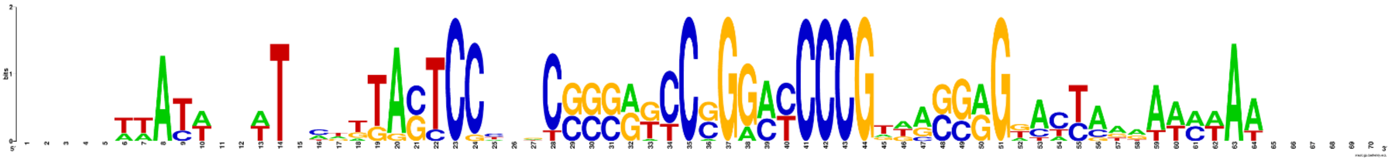

*Saccharomyces uvarum* strain CBS 395 (35 repeats)

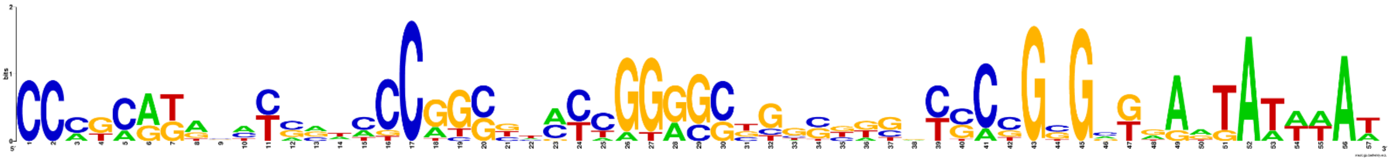

*Nakaseomyces bacillisporus* (91 repeats)

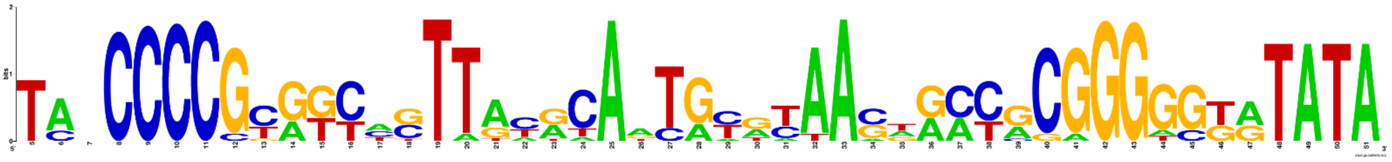

*Saccharomyces bayanus* strain CBS 380 (33 repeats)

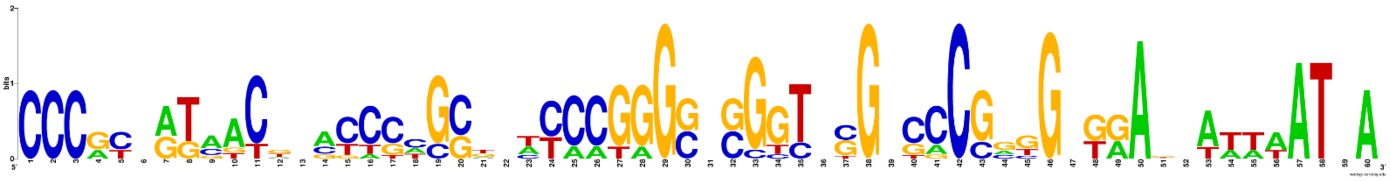

*Saccharomyces mikatae* strain CBS 8839 (19 repeats)

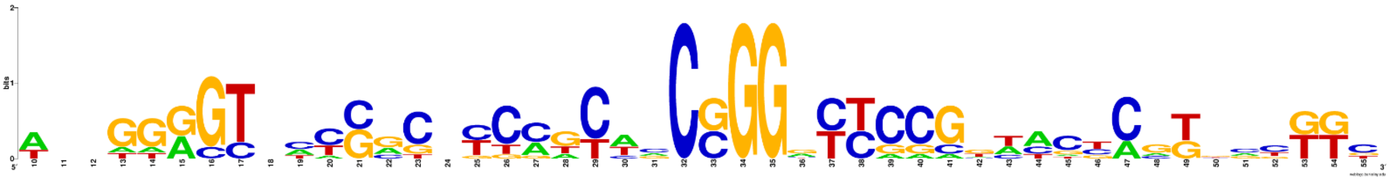

*Saccharomyces paradoxus* (9 repeats)

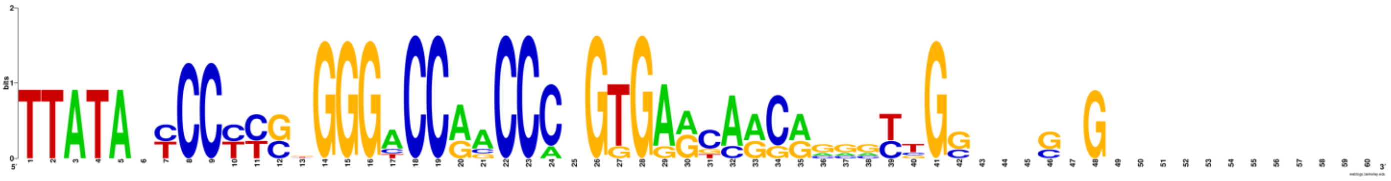

*Saccharomyces eubayanus* strain CBS12357 (32 repeats)

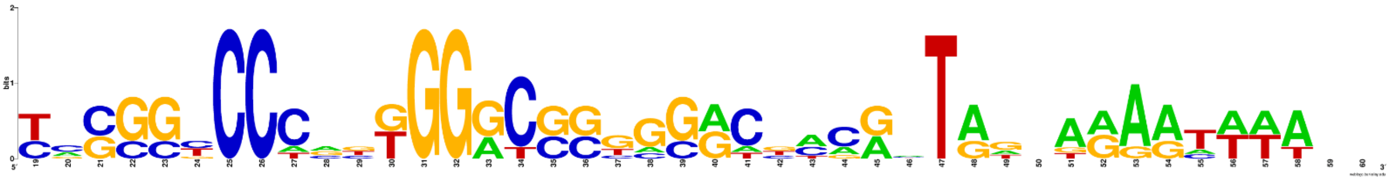

*Saccharomyces cariocanus* strain CBS 7994 (32 repeats)

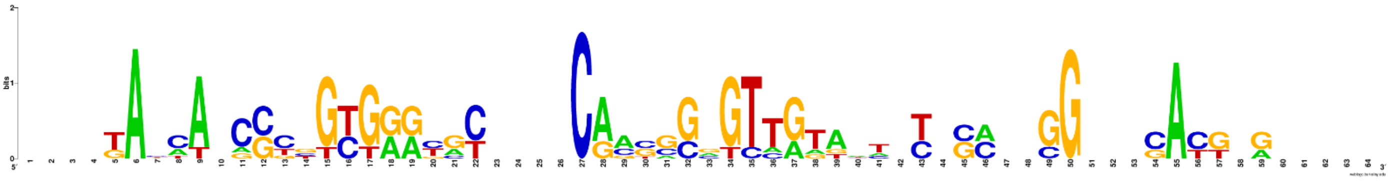

*Kazachstania sinensis* culture NRRLY-27222 (10 repeats)

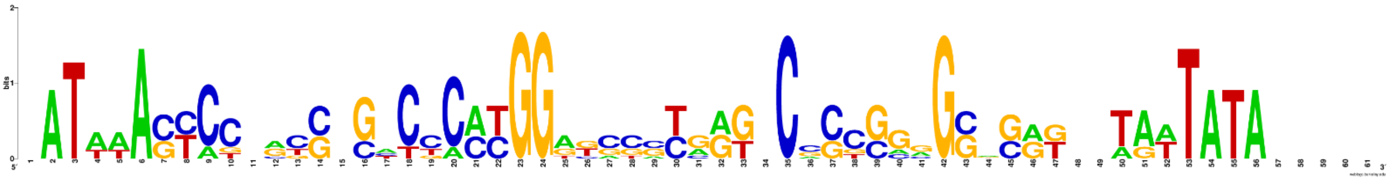

*Kazachstania unispora* strain Y-1556 (9 repeats)

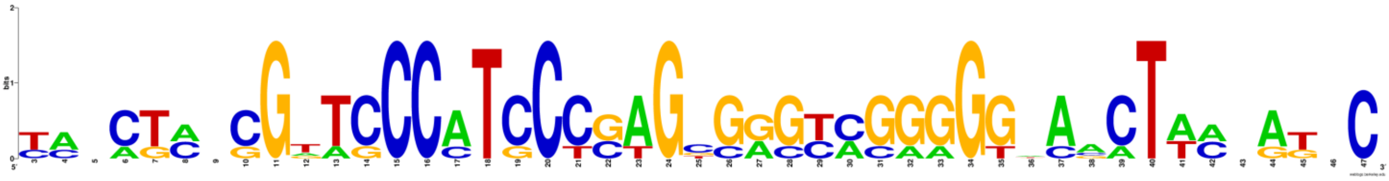

*Saccharomyces arboricola* strain NRRL Y-63701 (9 repeats)

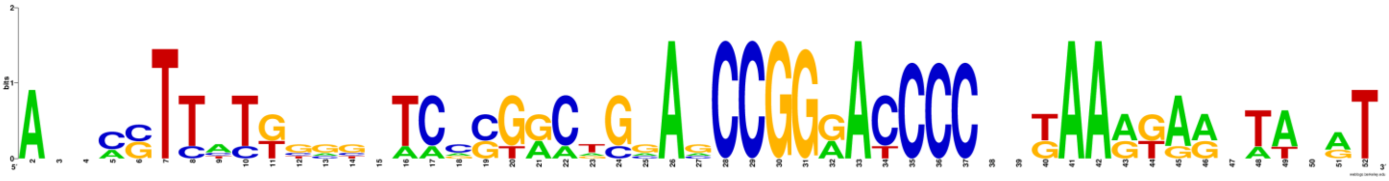

*Saccharomyces kudriavzevii* strain CR85 (60 repeats)

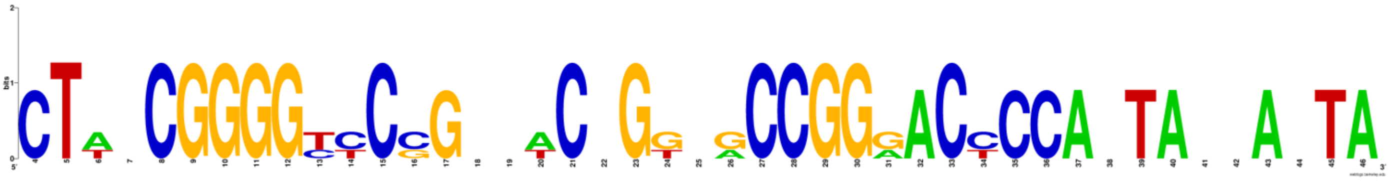

### Sporopachydermia clade

*Sporopachydermia lactativora* (226 repeats)

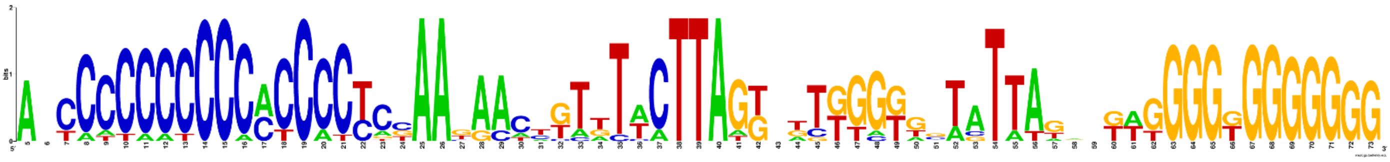

### Saccharomycetaceae TZ clade

*Torulaspora franciscae* strain Y6686 (44 repeats)

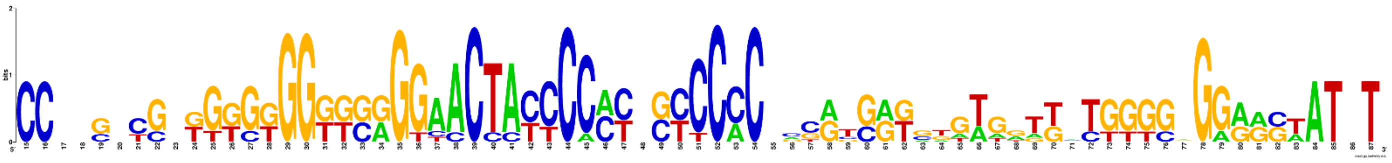

*Kluyveromyces lactis* strain GG799 (25 repeats)

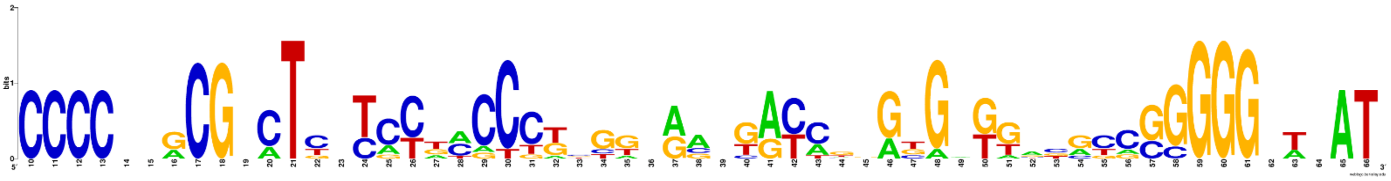

*Torulaspora globosa* strain CBS 2947 (13 repeats)

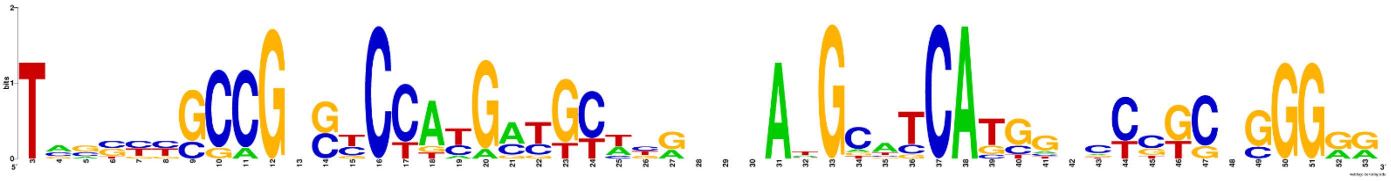

*Torulaspora franciscae* strain Y17532 (41 repeats)

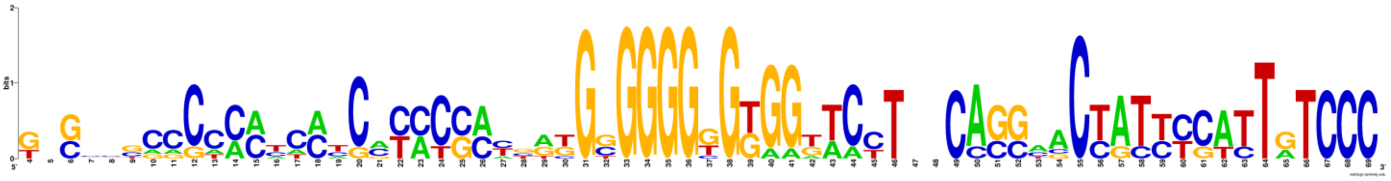

*Kluyveromyces lactis* (25 repeats)

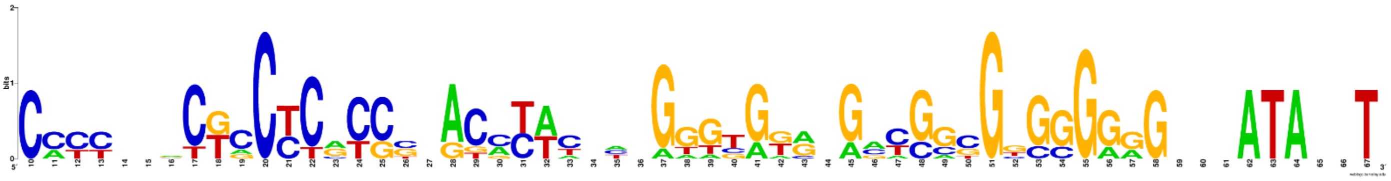

*Torulaspora delbrueckii* strain CBS 3003 (8 repeats)

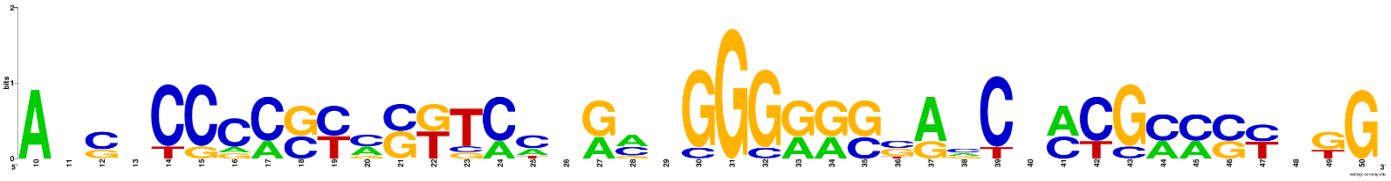

*Torulaspora delbrueckii* strain CBS 1146 (9 repeats)

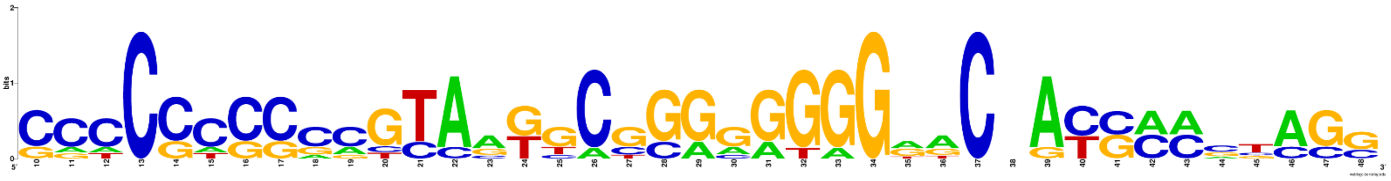

*Lachancea kluyveri* (2 repeats)

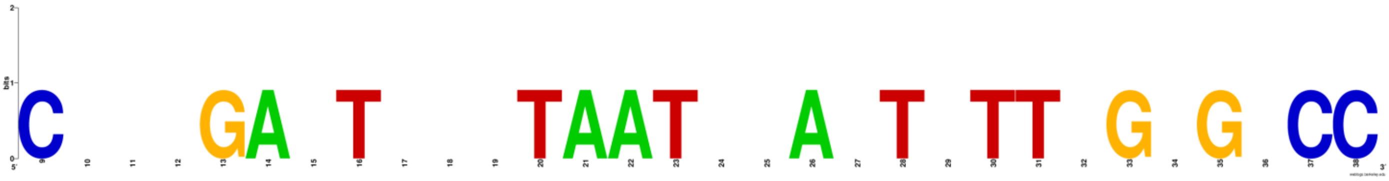

*Lachancea kluyveri* NCYC543 (3 repeats)

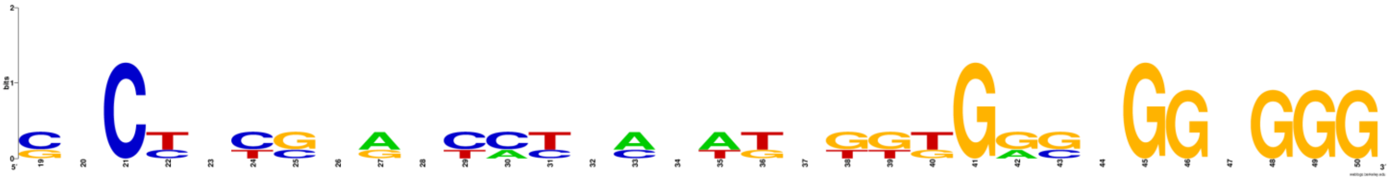

*Lachancea kluyveri* CBS5828 (4 repeats)

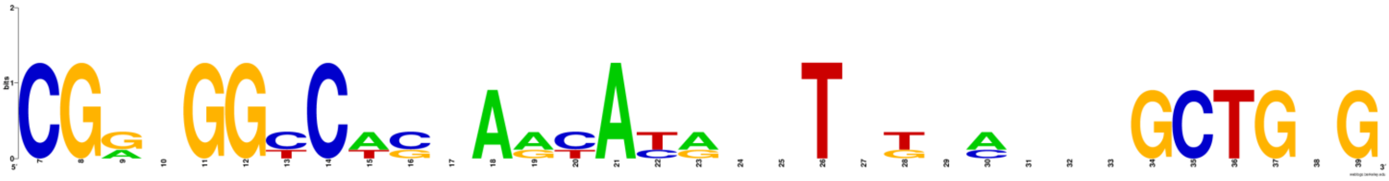

*Torulaspora delbrueckii* strain COFT1 (3 repeats)

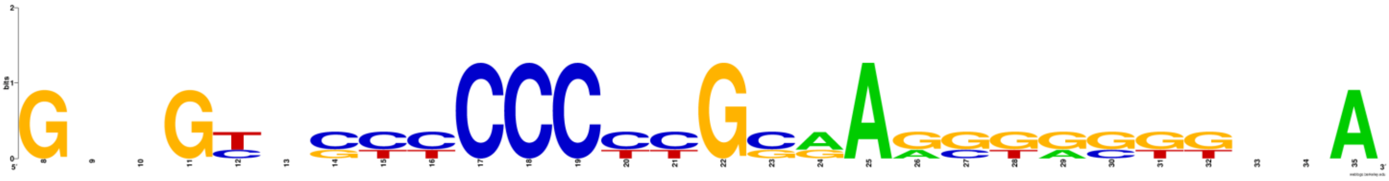

### Phaffomycetaceae clade

*Barnettozyma californica* strain CBS 252 (7 repeat)

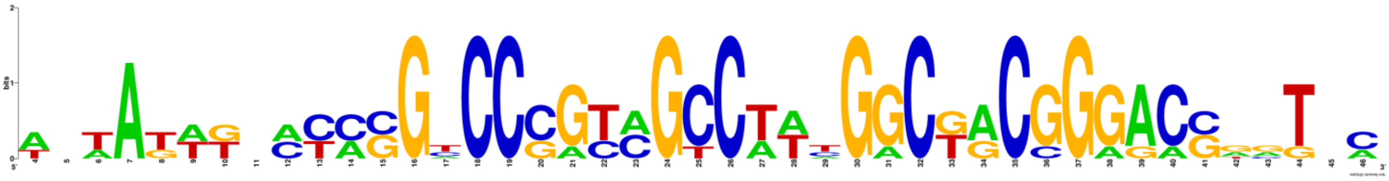
